# Development of an AAV Vector System for Highly Specific and Efficient Gene Expression in Microglia

**DOI:** 10.1101/2024.11.05.622037

**Authors:** Ryo Aoki, Ayumu Konno, Nobutake Hosoi, Hayato Kawabata, Hirokazu Hirai

## Abstract

Microglia play a critical role in diseases such as Alzheimer’s and stroke, making them a significant target for therapeutic intervention. However, due to their immune functions in detecting and combating viral invasion, efficient gene delivery to microglia remains challenging. We achieved specific and efficient gene delivery to microglia using an adeno-associated virus (AAV) vector designed for this purpose. This microglia-targeting AAV vector includes the mouse microglia/macrophage-specific ionized calcium-binding adaptor molecule 1 (mIba1) promoter, green fluorescent protein (GFP), microRNA target sequences (miR.Ts), woodchuck hepatitis virus posttranscriptional regulatory element (WPRE), and polyadenylation (polyA) signal, positioned between inverted terminal repeats. When miR.Ts were placed downstream of WPRE (between WPRE and polyA), gene expression occurred not only in microglia but also in a substantial number of neurons. However, when miR.Ts were positioned upstream of WPRE (between GFP and WPRE) or on both sides of WPRE, neuronal expression was significantly suppressed, resulting in selective GFP expression in microglia. Notably, positioning miR.Ts on both sides of WPRE achieved over 90% specificity and more than 60% efficiency in transgene expression in microglia three weeks after viral administration. This vector also enabled GCaMP expression in microglia, facilitating real-time monitoring of calcium dynamics and microglial process activity in the cortex. Additionally, intravenous administration of this vector with the blood-brain barrier-penetrant AAV-9P31 capsid variant resulted in extensive GFP expression selectively in microglia throughout the brain. These findings establish this AAV vector system as a robust tool for long-term, specific, and efficient gene expression in microglia.

## Introduction

Microglia, the resident immune cells of the central nervous system (CNS), originate from progenitor cells in the fetal yolk sac and play essential roles in monitoring and regulating neuronal activity.^1^ By extending their processes, microglia make dynamic contacts with synapses and axons, contributing to the maintenance of neuronal function.^2,3^ In pathological conditions, microglia become activated, migrating to lesions,^4^ phagocytosing damaged cells,^5,6^ and releasing various humoral factors.^7,8^ Microglial activation has been implicated in the pathogenesis of several CNS diseases, including Alzheimer’s disease,^9,10^ and multiple sclerosis,^11,12^ making these cells attractive therapeutic targets.

Given their critical role in brain homeostasis and disease, microglia have become important targets for genetic manipulation. However, selectively expressing genes in microglia using viral vectors has proven challenging, as microglia are involved in antiviral defense mechanisms within the CNS. In 2013, Jakobsson and colleagues successfully achieved selective transgene expression in microglia by utilizing lentiviral vectors containing the phosphoglycerate kinase (PGK) promoter and microRNA-9-target (miR-9.T) sequences.^13^ Because miR-9 is highly expressed in neurons and astrocytes but not in microglia, the presence of miR-9.T led to degradation of transgene mRNA in non-microglial cells. As a result, selective gene expression was achieved in microglia, with over 70% of transgene-expressing cells being microglia in the rat striatum. However, when the PGK promoter was replaced by the stronger cytomegalovirus (CMV) promoter, transgene expression leaked into neurons and astrocytes.^14^

To address these issues, we previously developed an adeno-associated virus serotype 9 (AAV9) vector that targets microglia. This vector combines a mouse-derived microglia/macrophage-specific ionized calcium-binding adaptor molecule 1 (mIba1) promoter with miR-9.T and miR-129-2-3p.T sequences.^15^ These microRNA targets, expressed in neurons but not in microglia, allowed for selective transgene expression in microglia within the striatum and cerebellum following brain parenchymal injection. However, while transgene expression in non-target neurons of the cerebral cortex was not prominent one week after injection, by three weeks, a substantial number of neurons exhibited strong transgene expression. This highlighted a major unresolved challenge in achieving microglia-specific expression in the cerebral cortex.^15^

Recent efforts to enhance microglial targeting have included the development of AAV capsid variants. Lin and colleagues identified two AAV9 capsid mutants, AAV-MG1.1 and AAV- MG1.2, through in vivo screening.^16^ Although these capsid mutants were able to transduce microglia, they also transduced neurons and astrocytes, limiting their specificity. Similarly, Young et al. developed AAV capsid mutants capable of crossing the blood-brain barrier (BBB) and efficiently transducing microglia.^17^ Although GFP expression was observed very efficiently in microglia, these mutants also induced GFP expression in neurons, oligodendrocytes, and astrocytes, highlighting the persistent challenge of achieving microglia-specific gene expression in the cerebral cortex.

Here, we aimed to overcome these limitations and, consequently, to develop AAV vectors with both high specificity and efficiency in transgene expression targeting cortical microglia. Our goal was to create a tool for long-term, microglia-specific gene expression in the cerebral cortex that could be used to study microglial function and serve as a platform for potential therapeutic interventions targeting microglia in neuropsychiatric disorders.

## Results

### The addition of a new miR-T (miR-708-5p.T×3) did not enhance neuron detargeting

In our previous study, we found that inserting quadruplet miR-9.T downstream of the woodchuck hepatitis virus posttranscriptional regulatory element (WPRE) in the AAV.mIba1.GFP.WPRE construct improved microglial specificity of GFP expression in the mouse cerebral cortex.^15^ Additionally, the inclusion of quadruplet miR-129-2-3p.T further enhanced microglial specificity.^15^ Since miR-708-5p is expressed in neurons but not in microglia,^18^ we investigated whether adding miR-708-5p.T to the AAV.mIba1.WPRE.GFP.miR- 9.T.miR-129-2-3p.T construct could improve neuron detargeting. For simplicity, the miR-9.T, miR-129-2-3p.T, and miR-708-5p.T sequences are abbreviated as “a”, “b”, and “c”, respectively (Fig. 1A).

**Figure 1.**
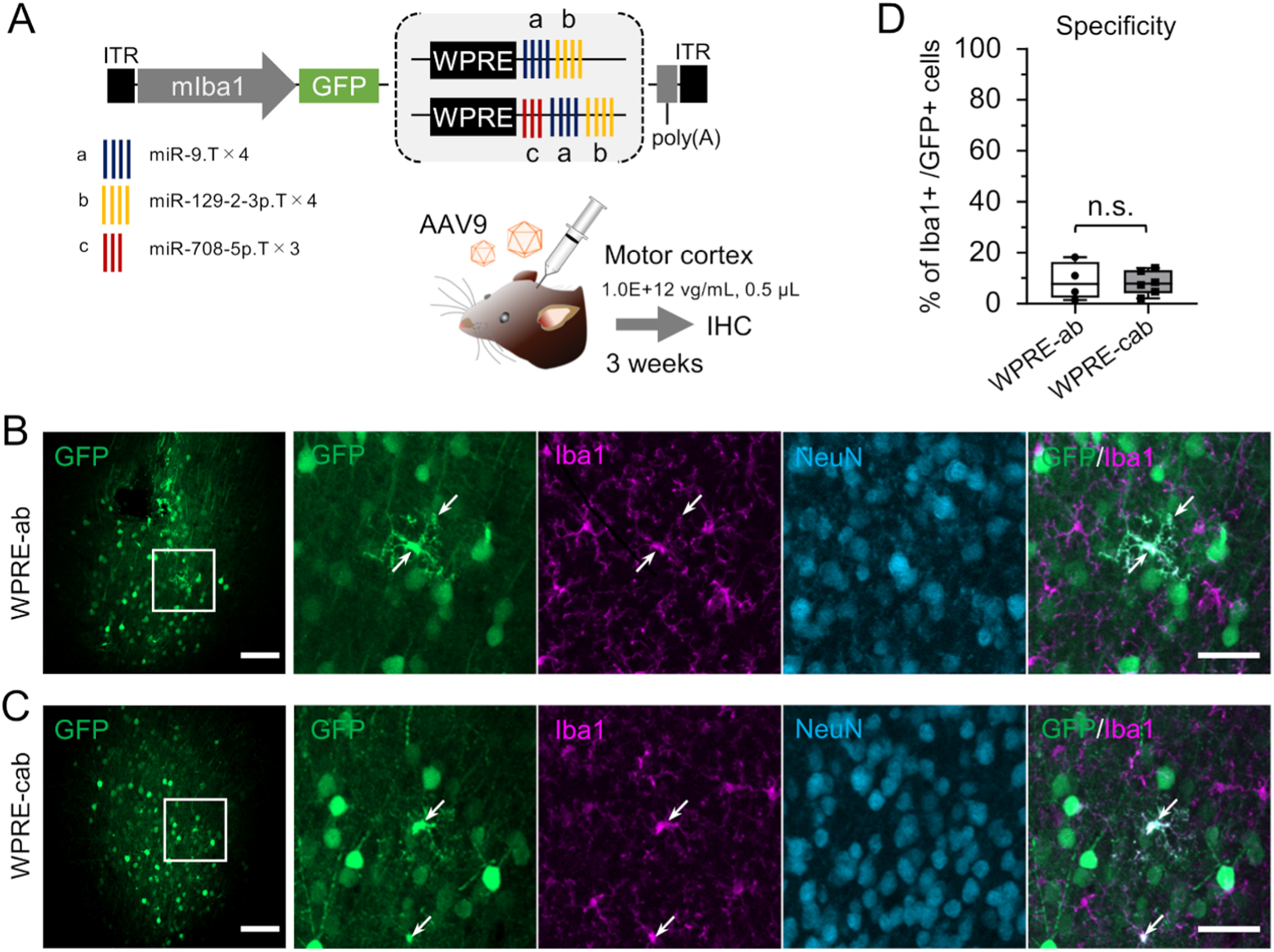
Addition of triplet miR-708-5p.T fails to enhance microglia detargeting. **(A)** Schematic depicting the AAV genome comprising microRNA target sequences: the quadruplet or triplet microRNA target sequences miR-9.T×4, miR-129-2-3p.T×4, and miR- 708-5p.T×3, abbreviated as “a”, “b”, and “c”, respectively. Triplet miR-708-5p.T was inserted between WPRE and the quadruplet miR-9.T in the genome of a previously reported microglia-targeting AAV harboring WPRE-ab. The genomes were packaged with the AAV9 capsid. Six- to eight-week-old C57BL/6J mice received an injection of either one of the AAV vectors (1.0E+12 vg/mL, 0.5 μL) into the motor cortex. Three weeks after injection, mice were euthanized, and cerebral sections were prepared and analyzed by immunohistochemistry (IHC). **(B, C)** Confocal laser-scanning microscopy of cerebral sections injected with a control AAV carrying WPRE-ab (B) and those injected with AAV carrying WPRE-cab (C). The sections were triple immunostained for GFP, Iba1, and NeuN. Arrows indicate GFP-expressing Iba1- positive microglia. Scale bars: 100 μm for the low magnification image on the left, and 20 μm for the enlarged images on the right. **(D)** Summarized graph showing the specificity of microglia transduction in the two mouse groups.. n.s., not significant by unpaired t-tests (n = 4 hemispheres for WPRE-ab, n = 6 hemispheres for WPRE-cab). The box-and-whisker plots depict the median (centerlines), 25th and 75th percentiles (bounds of the box), and minimum/maximum values (whiskers).

We injected AAV.mIba1.GFP.WPRE-cab (1.0E+12 vg/mL, 0.5 μL) or AAV.mIba1.GFP.WPRE-ab into the motor cortex of mice (n = 6 for WPRE-cab, n = 4 for WPRE-ab) and analyzed brain sections three weeks later. Immunohistochemistry revealed no significant differences in the number of GFP-expressing neurons between the WPRE-cab and WPRE-ab groups (Fig. 1B, C). Quantitative analysis confirmed that the addition of miR-708- 5p.T did not significantly improve neuron detargeting, as the microglial specificity of GFP expression was similar between WPRE-cab (8.2 ± 4.2%, n = 6 hemispheres) and WPRE-ab (8.8 ± 6.4%, n = 4 hemispheres) (Fig. 1D).

### Placing microRNA target sequences upstream of WPRE significantly enhances neuron detargeting

During AAV transduction in neurons, the viral genome is transported to the nucleus, where mRNA is transcribed. It has been shown that miR.T-containing mRNA is cleaved between the 9th and 10th base pairs downstream from the 5’ side of the miR.T sequence.^19,20^ When miR.T sequences are positioned downstream of WPRE (AAV.mIba1.WPRE-ab), the mRNA is cleaved at the site downstream of WPRE, resulting in an mRNA consisting of GFP and WPRE (Fig. S1A, B). WPRE stabilizes the mRNA, which enhances protein expression levels, independent of the transgene or promoter.^21^ Consequently, it is hypothesized that even though the resulting GFP-WPRE mRNA lacks a polyadenylation [poly(A)] signal, it is still translated into GFP protein due to the stabilizing effects of WPRE. In contrast, when the miR.T sequence is inserted between GFP and WPRE, the mRNA is cleaved between these two elements, resulting in an mRNA consisting only of GFP (Fig. S1C). However, this truncated GFP mRNA lacks both WPRE and the poly(A) signal, leading to degradation, and it is expected that no GFP protein will be produced from this mRNA.

To confirm whether GFP protein expression occurs in the presence of WPRE without poly(A), but not in its absence, we prepared AAVs with the following sequences in the AAV genome: mIba1.GFP.WPRE.poly(A), mIba1.GFP.WPRE-ab.poly(A), mIba1.GFP.WPRE, and mIba1.GFP (Fig. S1D). These AAVs were injected into the motor cortex of mice (1.0E+12 vg/mL, 1.0 μL), and GFP fluorescence was observed three weeks post-injection. As a result, strong GFP expression was observed in mIba1.GFP.WPRE.poly(A), while weak but comparable GFP expression was detected in mIba1.GFP.WPRE-ab.poly(A) and mIba1.GFP.WPRE (Fig.S1E-G). It was hypothesized that GFP.WPRE-ab.poly(A) mRNA is cleaved at the ab (miR.T) site, producing GFP-WPRE mRNA. Additionally, it was found that GFP protein production was nearly absent when both WPRE and poly(A) were missing (Fig. S1H). These results suggested that placing miR.T between GFP and WPRE enhances the suppression of GFP protein expression due to mRNA cleavage between GFP and WPRE (Fig. S1C).

To test this, we created two AAV vectors: AAV.mIba1.GFP.ab-WPRE and AAV.mIba1.GFP.ab-WPRE-ab, where the miR.T sequences were positioned upstream and on both sides of WPRE, respectively (Fig. 2A). These vectors were injected into the motor cortex of mice.

**Figure 2.**
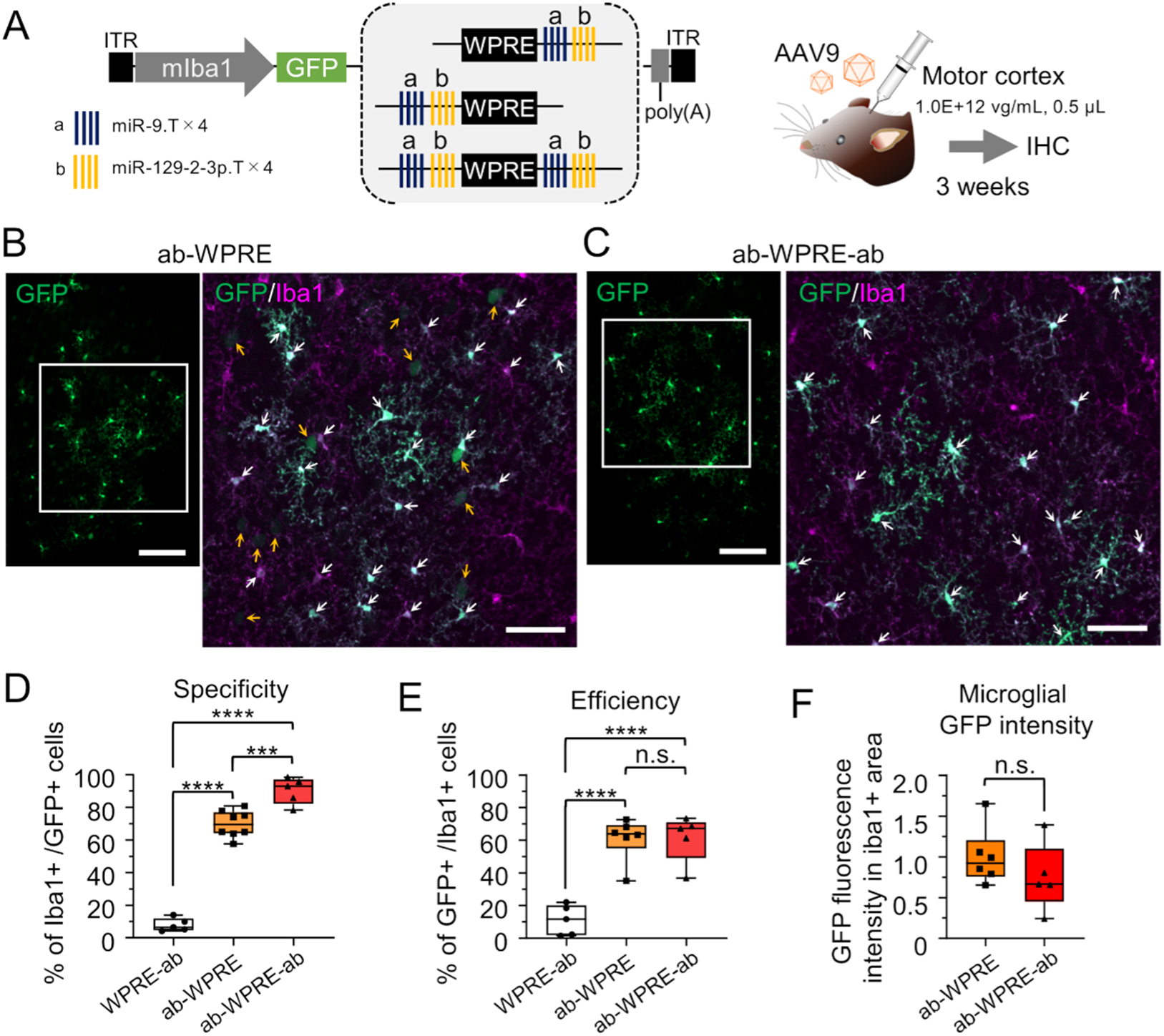
Significant enhancement of specificity and efficiency of GFP expression in microglia by placing the miR.Ts upstream of WPRE. **(A)** Schematic depicting the AAV genome with microRNA target sequences placed on the 3’ side (control), 5’ side, or both sides of WPRE. Mice received an injection of either one of the AAV9 vectors (1.0E+12 vg/mL, 0.5 μL) into the motor cortex and were euthanized for immunohistochemistry (IHC) three weeks post-injection. **(B, C)** Confocal microscopy of cerebral sections immunostained from mice injected with AAV.mIba1.GFP.ab-WPRE (ab-WPRE) (B) and those injected with AAV.mIba1.GFP.ab-WPRE-ab (ab-WPRE-ab) (C). For the left and right panels, the left side shows low magnification images of GFP immunostaining, while the right side shows high magnification images of GFP and Iba1 immunostaining, which are enlarged views of the boxed areas in the left images. White and yellow arrows indicate transduced microglia and neurons, respectively. Scale bars: 100 μm (left, low magnification) and 50 μm (right, enlarged images). **(D, E)** Summarized graphs showing specificity (n = 5 hemispheres for WPRE-ab, n = 8 hemispheres for ab-WPRE, n = 5 hemispheres for ab- WPRE-ab) (D) and efficiency (n = 5 hemispheres for WPRE-ab, n = 6 hemispheres for ab- WPRE, n = 5 hemispheres for ab-WPRE-ab) (E) of microglia transduction in the three mouse groups. Efficiency of microglia transduction was calculated as the number of GFP- and Iba1- double-positive cells divided by the number of Iba1-immunolabeled microglia within a 320 μm × 320 μm area. ***p < 0001, ****p < 0.0001, n.s., not significant by Bonferroni’s multiple comparisons test following one-way ANOVA. **(F)** A graph comparing GFP fluorescence values in the Iba1+ area of mice injected with AAV carrying ab-WPRE or AAV carrying ab-WPRE-ab. The total GFP fluorescence intensity in a 320 μm x 320 μm Iba1+ area was measured, setting the value for AAV carrying ab-WPRE at 1. n.s., not significant by unpaired t-tests. (D-F) The box-and-whisker plots depict the median (centerlines), 25th and 75th percentiles (bounds of the box), and minimum/maximum values (whiskers). See also Figure S1-S3.

Immunohistochemistry revealed that GFP expression was highly specific to microglia in both constructs. Quantitative analysis showed that the microglial specificity of GFP expression was significantly higher in AAV vectors with miR.T sequences upstream of WPRE (ab-WPRE: 69.9 ± 7.6%, n = 8 hemispheres) compared to the previously reported WPRE-ab (7.9 ± 3.6%, n = 5 hemispheres) (****p < 0.0001 by Bonferroni’s multiple comparisons test following one-way ANOVA) (Fig. 2D). Additionally, the efficiency of GFP expression in microglia also significantly increased, with 61.0 ± 12.1% of microglia expressing GFP in the ab-WPRE condition (n = 6 hemispheres), 61.5 ± 12.9% for ab-WPRE-ab (n = 5 hemispheres), and 11.1 ± 8.3% for WPRE-ab (n = 5 hemispheres) (****p < 0.0001 vs. WPRE-ab by Bonferroni’s multiple comparisons test following one-way ANOVA) (Fig. 2E).

Notably, placing miR.T sequences on both sides of WPRE (ab-WPRE-ab) further enhanced microglial specificity compared to miR.T sequences positioned only upstream of WPRE. Microglial specificity reached 90.3 ± 7.3% with ab-WPRE-ab (n = 5 hemispheres), compared to 69.9 ± 7.6% with ab-WPRE (n = 8 hemispheres) (***p < 0.001 by Bonferroni’s multiple comparisons test following one-way ANOVA) (Fig. 2D). There were no significant differences between ab-WPRE and ab-WPRE-ab in the efficiency of microglia transduction or the GFP fluorescence intensity in transduced microglia (p > 0.9999 by Bonferroni’s multiple comparisons test and p = 0.3122 by unpaired t-test, respectively) (Fig. 2E, F).

The expression levels of GFP in microglia were thought to be higher in AAVs with miR.T sequences upstream of WPRE compared to AAVs with miR.T sequences only downstream of WPRE. This is because GFP fluorescence was clearly observable without immunohistochemistry in sections treated with AAVs containing miR.T sequences upstream of WPRE (ab-WPRE and ab-WPRE-ab), whereas it was scarcely detectable in sections treated with AAVs containing miR.T sequences only downstream of WPRE (WPRE-ab) (Fig. S2).

The higher microglial specificity of ab-WPRE-ab compared to ab-WPRE is unlikely to be due to the presence of two copies of miR.T, as placing two copies of miR.T upstream of WPRE (abab-WPRE) did not improve neuron detargeting (Fig. S3).

### The Iba1 promoter is indispensable for transgene expression in microglia

Given the significant enhancement in microglial specificity with miR.T sequences placed upstream of WPRE, we next investigated the necessity of the mIba1 promoter for microglia-specific gene expression. We replaced the mIba1 promoter with the ubiquitously active cytomegalovirus early enhancer/chicken β-actin (CAG) promoter in AAV vectors expressing GFP-ab-WPRE-ab-poly(A) and injected these into the cerebral cortex of mice (n = 4 hemispheres) (Fig. 3A). Surprisingly, we found that GFP was not expressed in microglia, but instead was predominantly expressed in oligodendrocytes and, to a lesser extent, astrocytes (Fig. 3B-D).

**Figure 3.**
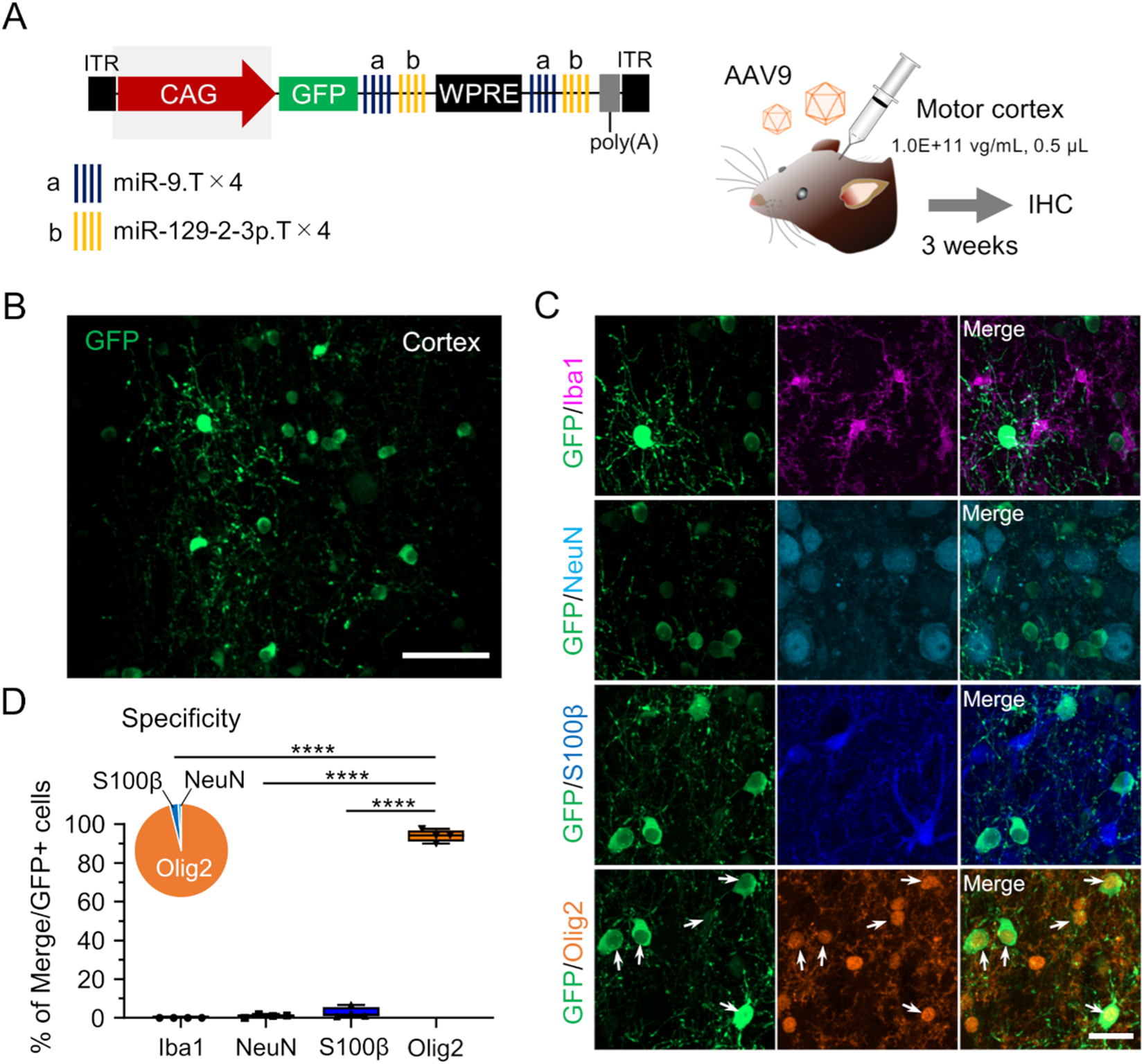
Oligodendrocyte-specific GFP expression by AAV.CAG.GFP.ab-WPRE-ab. **(A)** Schematic depicting the AAV genome comprising microRNA target sequences on both sides of WPRE and expressing GFP under the CAG promoter. Mice received an injection of the AAV9 vectors (1.0E+11 vg/mL, 0.5 μL) into the motor cortex. Mice were euthanized three weeks post-injection and analyzed by immunohistochemistry (IHC). **(B, C)** Low-magnification fluorescent images immunolabeled for GFP (B) and enlarged images of the transduced areas immunostained for GFP, Iba1, NeuN, S100β, and Olig2 (C). Arrows indicate Olig2- immunolabeled GFP-expressing oligodendrocytes. Scale bars: 100 μm (B) and 20 μm (C). **(D)** Box-and-whisker graph and pie chart showing the percentage of each cell type among total GFP-expressing cells (n = 4 hemispheres). Note that almost all GFP-expressing cells are Olig2-labeled oligodendrocytes. ****p < 0.0001 by Bonferroni’s multiple comparisons test following one-way ANOVA. The box-and-whisker plots depict the median (centerlines), 25th and 75th percentiles (bounds of the box), and minimum/maximum values (whiskers).

This unexpected result indicates that while the miR-9.T and miR-129-2-3p.T sequences are sufficient to suppress transgene expression in neurons and astrocytes, oligodendrocytes, which lack or have low expression of these microRNAs,^22^ can express the transgene under the control of the CAG promoter. Furthermore, the lack of GFP expression in microglia suggests that the CAG promoter is not active in these cells, underscoring the importance of the mIba1 promoter for driving gene expression specifically in microglia.

### High microglial specificity of AAV with miR.T on both sides of WPRE in the striatum and cerebellum

To investigate whether the optimized AAV vector (AAV.mIba1.GFP.ab-WPRE-ab) could achieve microglia-specific transgene expression in other brain regions, we injected this vector into the striatum (n = 7 hemispheres) and cerebellum (n = 4 mice) of mice (Fig. 4A). Three weeks post-injection, we observed highly specific GFP expression in microglia in both brain regions (Fig. 4B, C). Compared with the control AAV.mIba1.GFP.WPRE-ab, the optimized AAV.mIba1.GFP.ab-WPRE-ab exhibited significantly higher specificity for microglia in the striatum (96.7 ± 2.3%, n = 7 hemispheres vs. 64.8 ± 20.0%, n = 8 hemispheres for control, **p< 0.01 by unpaired t-test) and almost comparable specificity for cerebellar microglia (91.9 ± 6.3%, n = 4 mice vs. 92.4 ± 9.0%, n = 4 mice for control) (Fig. 4D, F).

**Figure 4.**
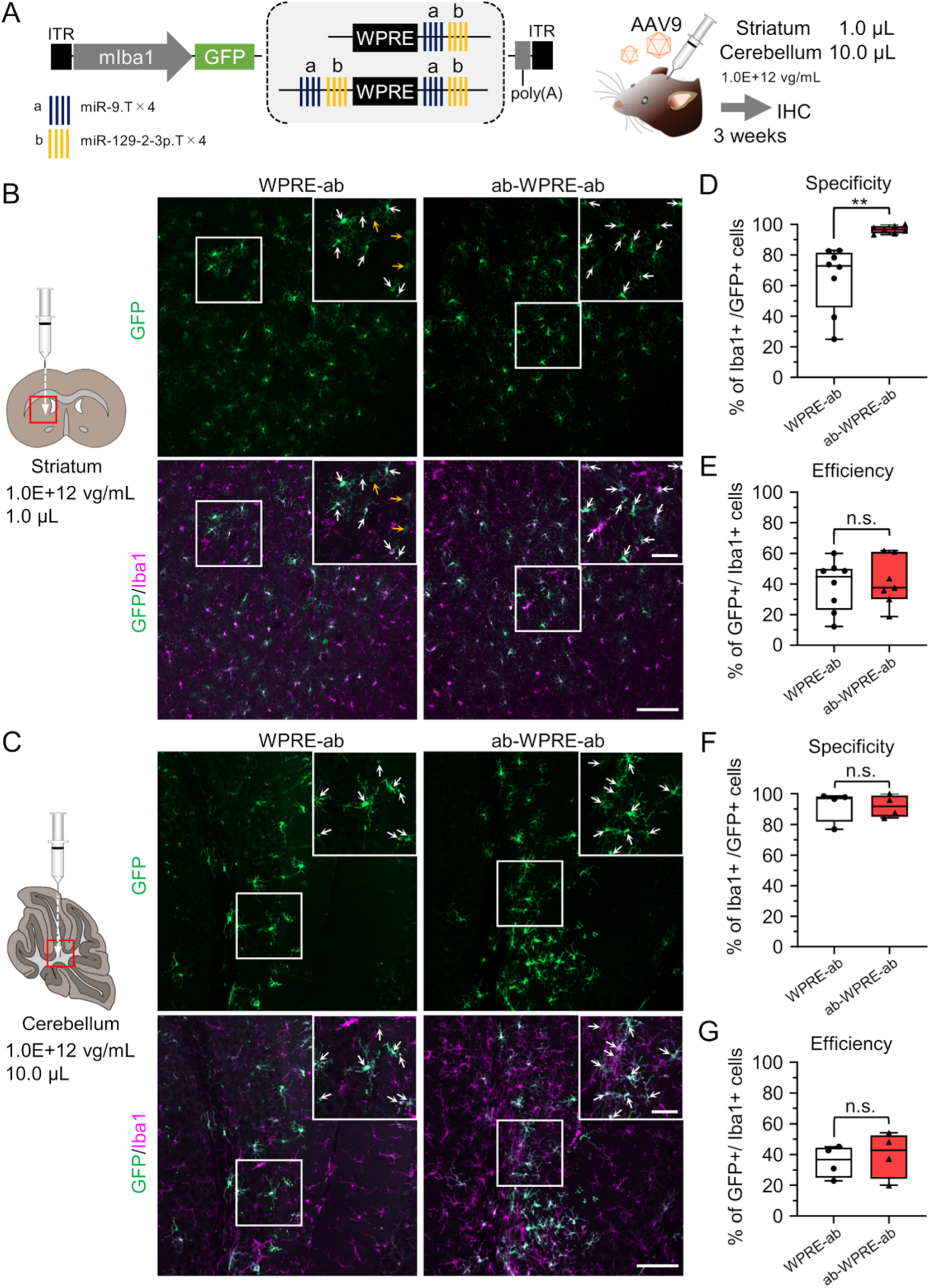
Microglia-specific GFP expression in the striatum and cerebellum by AAV.mIba1.GFP.ab-WPRE-ab. **(A)** AAV9.mIba1 carrying ab-WPRE-ab was injected into the striatum (1.0E+12 vg/mL, 1.0 µL) or cerebellum (1.0E+12 vg/mL, 10 µL) of C57BL/6J mice. Three weeks after the injection, the mice were euthanized and analyzed by immunohistochemistry (IHC). **(B, C)** Confocal microscopy of striatal (B) and cerebellar (C) sections. The left and right panels show sections from mice injected with AAV.mIba1.WPRE-ab (left) and AAV.mIba1.ab-WPRE-ab (right). Boxed areas are expanded in the upper right corner of each panel, where white and yellow arrows indicate GFP-expressing microglia and neurons, respectively. Scale bars: 100 μm (bottom right) and 40 μm (inset). **(D–G)** Summarized graphs showing the specificity (D, F) and efficiency (E, G) of GFP expression in microglia in the striatum (D, E; n = 8 hemispheres for WPRE-ab, n = 7 hemispheres for ab-WPRE-ab) and cerebellum (F, G; n = 4 hemispheres for WPRE-ab, n = 4 hemispheres for ab-WPRE-ab). **p < 0.01 by unpaired t-test. n.s., not significant. The box-and-whisker plots depict the median (centerlines), 25th and 75th percentiles (bounds of the box), and minimum/maximum values (whiskers).

The efficiency of GFP expression in microglia was comparable between the two AAV vectors in both the striatum and cerebellum (Str: 38.8 ± 15.4%, n = 8 hemispheres; Cbl: 35.4 ± 9.0%, n = 4 mice for WPRE-ab and Str: 41.2 ± 14.6%, n = 7 hemispheres; Cb: 39.9 ± 13.0%, n = 4 mice for ab-WPRE-ab) (Fig. 4E, G). Therefore, it is suggested that ab-WPRE-ab is capable of microglia-specific expression in other brain regions. The lack of a significant difference in microglial specificity between WPRE-ab and ab-WPRE-ab in the cerebellum may be because the mIba1 promoter already achieves high specificity without the need for miR.T in the cerebellum.^15^

### Sustained microglia-specific GFP expression two months post-injection

To evaluate the long-term persistence of microglia-specific gene expression, we injected AAV.mIba1.GFP.ab-WPRE-ab into the cerebral cortex and analyzed the brains two months post-injection (n = 6 hemispheres) (Fig. 5A). Although the frequency of GFP-positive non-microglial cells increased over time, GFP-expressing microglia still accounted for approximately 74.2 ± 11.1% of the total GFP-expressing cells (n = 6 hemispheres) (Fig. 5B- D). This suggests that while some leakage into non-microglial cells occurs over time, the vector retains high specificity for microglia over an extended period.

**Figure 5.**
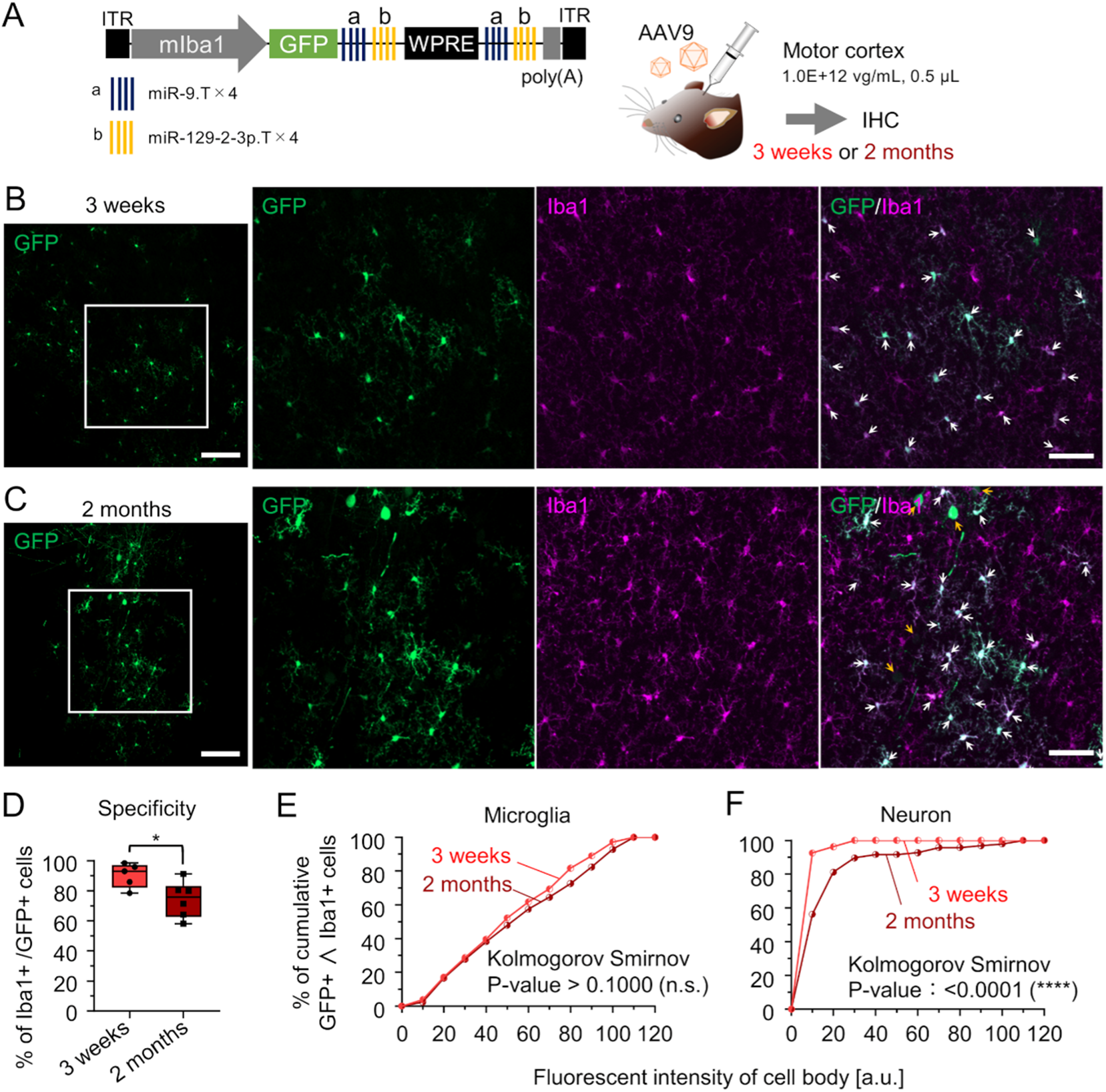
Maintenance of GFP expression specificity in microglia two months after AAV injection. **(A)** AAV.mIba1.GFP.ab-WPRE-ab (1.0E+12 vg/mL, 0.5 μL) was injected into the motor cortex of C57BL/6J mice. The mice were euthanized three weeks or two months after the injection and analyzed by immunohistochemistry (IHC). **(B, C)** Confocal microscopy of cerebral sections three weeks (B) and two months (C) post-injection. The two panels on the left are low-magnification images, while the panels on the right are magnified images of the boxed areas in the left panels. White and yellow arrows indicate GFP-expressing microglia and non-microglial cells, respectively. Scale bars: 100 μm (left) and 50 μm (right). **(D)** Graph showing the specificity of GFP expression in microglia three weeks (n = 5 hemispheres) or two months (n = 6 hemispheres) after AAV injection. The data for the 3-week time point is the same as in Fig. 2D. *p < 0.05 by unpaired t-test. **(E, F)** Cumulative plot of GFP fluorescence intensity in the cell bodies of GFP- and Iba1-double-positive microglia (E) or in the cell bodies of GFP-positive and Iba1-negative neurons (F). Red and dark red symbols indicate results obtained from mice three weeks after injection (285 microglial cell bodies (E) and 27 neuronal cell bodies (F) from 5 hemispheres) and results obtained from mice two months after injection (296 microglial cell bodies (E) and 96 neuronal cell bodies (F) from 6 hemispheres), respectively. p > 0.1 (n.s., not significant) for (E) and ****p < 0.0001 for (F) by Kolmogorov- Smirnov test.

The intensity of GFP fluorescence in microglia did not significantly change between the three-week and two-month time points, indicating that transgene expression levels remained stable in microglia over time (p > 0.1 by Kolmogorov Smirnov test) (Fig. 5E). In contrast, GFP fluorescence intensity in neurons increased significantly (****p > 0.0001 by Kolmogorov Smirnov test), suggesting that the gradual reduction in microglial specificity was due to increased transgene expression in non-target neurons (Fig. 5F).

### Application of our optimized microglia-selective gene expression method to physiological experiments in cortical microglia

Next, we examined whether our optimized microglia-selective gene expression method could be applied to physiological experiments involving microglia in the motor cortex. First, we attempted to measure Ca²⁺ signals through AAV-mediated expression of the genetically encoded fluorescent calcium indicator, jGCaMP8s, in cortical microglia of the primary and secondary motor areas.^23^ Three to four weeks after AAV injection into the motor cortex, we performed confocal live Ca²⁺ imaging in acute cerebral slices, where GCaMP-positive microglia were observed using our updated method (Fig. 6A).

**Figure 6.**
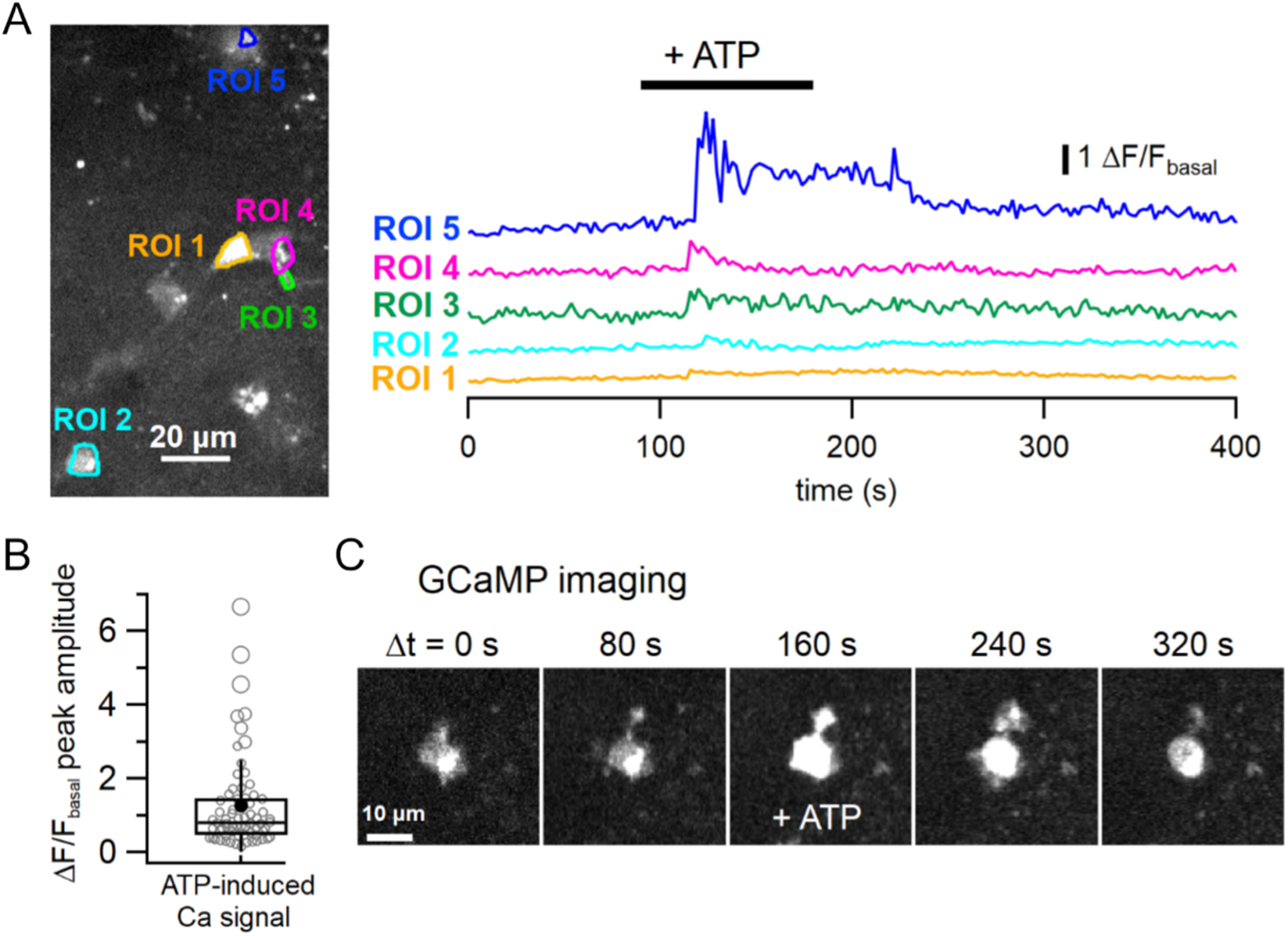
Measurement of Ca²⁺ signals in cortical microglia expressing the genetically encoded Ca²⁺ indicator jGCaMP8s using our microglia-selective AAV-mediated gene expression method. **(A)** The left panel displays an averaged confocal image of virally expressed jGCaMP8s signals in the motor cortex of an acute cerebral slice. Regions of interest (ROIs 1–5) were placed on microglial compartments. The right panel shows Ca²⁺ signal traces estimated from the ROIs in the left panel. Bath application of 100 µM ATP (for 90 s, indicated by the black bar) induced Ca²⁺ transients in the transduced microglia. **(B)** A box-and-whisker plot showing the peak amplitude of quantified Ca²⁺ signals induced by bath-applied ATP (ΔF/Fbasal; see Methods) in microglial compartments. Open circles represent individual data points, while the horizontal line and the box represent the median value and interquartile range, respectively (n = 68 cellular compartments from eight cerebral slices of five mice). The error bars extend one standard deviation above and below the mean (filled circle). **(C)** Time-lapse GCaMP images (single focal plane) capturing both the movement of microglial processes and ATP-induced Ca²⁺ increases in the microglia. See also Video S1.

Bath application of ATP (100 µM) induced a Ca²⁺ increase in cellular compartments, including microglial cell bodies and processes, with a variable delay (Fig. 6A; Video S1). This delay is likely due to the time required for solution exchange and the microglial response. The mean peak amplitude of ATP-induced Ca²⁺ signal changes (ΔF/Fbasal; see Methods) was 1.26 ± 0.15 (Fig. 6B; n = 68 cellular compartments from eight cerebral slices of five mice). These results align with typical microglial Ca²⁺ dynamics, as microglia express purinergic receptors and exhibit ATP-induced Ca²⁺ responses within tens of seconds.^24^ Additionally, some GCaMP-positive cells exhibited process movement and ATP-induced process extension (Fig. 6C; Video S1), which are hallmark features of microglia.^25^

Since the GCaMP signals in Fig. 6C reflect both the movement of the processes and changes in Ca²⁺ concentration within the processes, it is difficult to discern whether the processes themselves are moving or only the Ca²⁺ concentration is changing. To more accurately capture dynamic morphological changes and process movements in microglia, we specifically expressed GFP in cerebral microglia using our method and performed live GFP imaging with confocal microscopy (Fig. 7). Most GFP-positive microglia exhibited clear basal morphological motility, although there was some variability among individual cells (Fig. 7A; Videos S2-5). In many cases, it was difficult to reliably capture the complete three-dimensional movement of a single microglial process within a single focal plane (Fig. 7A, B, upper panels; Video S6). To overcome this, we acquired z-stack images (typically more than 25 images), covering the entire structure of a single microglia at each time point, and created time-lapse two-dimensional images from the maximum intensity projections of the z-stacks (Fig. 7A, B, lower panels).

**Figure 7.**
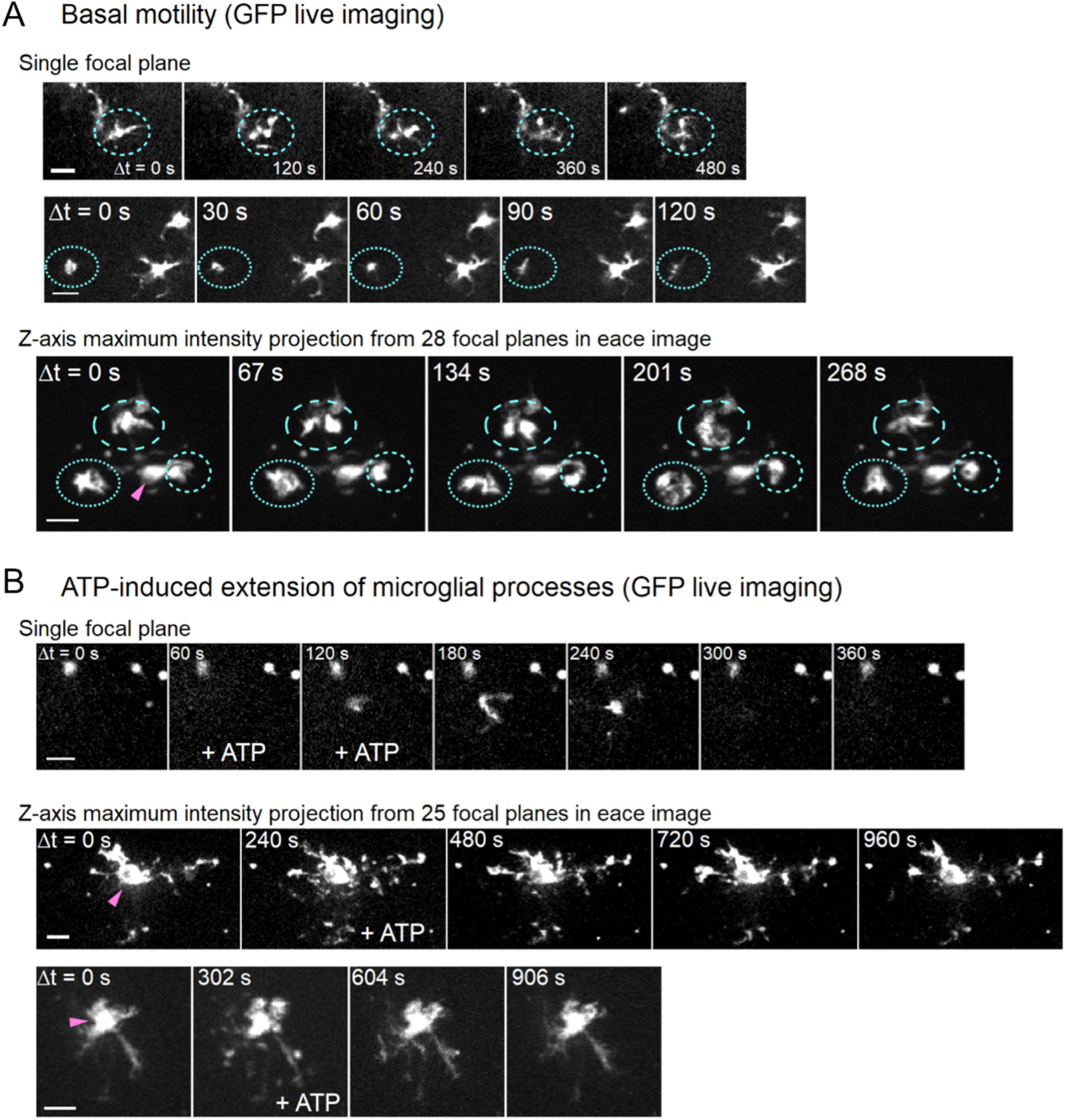
Monitoring microglial motility in the mouse motor cortex using our microglia- selective AAV-mediated gene expression method. **(A, B)** Time-lapse GFP fluorescent images with time points indicated, showing basal movements of microglial processes in (A) and ATP-induced process extension in (B). Frames during the bath application of 100 µM ATP (3–4.5 minutes) are labeled as ‘+ ATP’. Blue circles highlight areas with pronounced basal motility, and magenta arrowheads indicate the putative cell bodies of microglia in maximum intensity projection images of z-stacks. Each row of time-lapse images in (A) and (B) represents different microglia. Scale bars: 10 μm. See also Video S2-S9.

Time-lapse 2D images from multiple focal planes showed that microglial cell bodies remained relatively static, while the processes exhibited dynamic motility (Fig. 7A, B; arrowheads; Videos S4-5 and S7-9). Bath application of ATP (100 µM) induced pronounced elongation of microglial processes after a delay (Fig. 7B; Videos S6-9), consistent with the typical morphological dynamics of microglia.^25,26^ These results suggest that our optimized microglia-selective expression method is effective for live imaging experiments investigating the morphological dynamics of microglia. Taken together, we conclude that our updated AAV-mediated, microglia-specific gene expression method is also applicable to physiological experiments.

### Successful microglial transgene expression by intravenous injection of AAV.mIba1 harboring ab-WPRE-ab

We next investigated whether our updated AAV vector could achieve microglia-specific gene expression through intravenous injection. To this end, we used five different blood-brain barrier (BBB)-penetrating capsid variants—PHP.B,^27^ PHP.eB,^28^ 9P31,^29^ and two different innate-MG [AAV(H) and AAV(Y)] —to package the AAV genome (Fig. S4A, B).^17^ Innate-MG capsid variants feature a seven-amino acid insertion, (HGTAASH) or (YAFGGEG), between amino acids 588 and 589 of the AAV9 capsid. These variants have been shown to deliver transgenes to microglia with up to 80% efficiency following intravenous injection.^17^ Mice received intravenous injection of one of these AAV vectors (2.0E+13 vg/mL, 100 μL). Three weeks post-injection, brain sections were analyzed by immunohistochemistry.

Fluorescence microscopy revealed that brain sections from mice injected with PHP.B, PHP.eB, and innate-MG vectors had few GFP-labeled cells, whereas sections from mice injected with the AAV-9P31 vector showed more GFP-expressing microglia, although the signal was faint (Fig. S4C-G). To enhance transgene expression and more clearly label microglia, we increased the injection dose of the AAV-9P31 vector to 6.8E+13 vg/mL (100 μL) and repeated the experiment (Fig. 8A).

**Figure 8.**
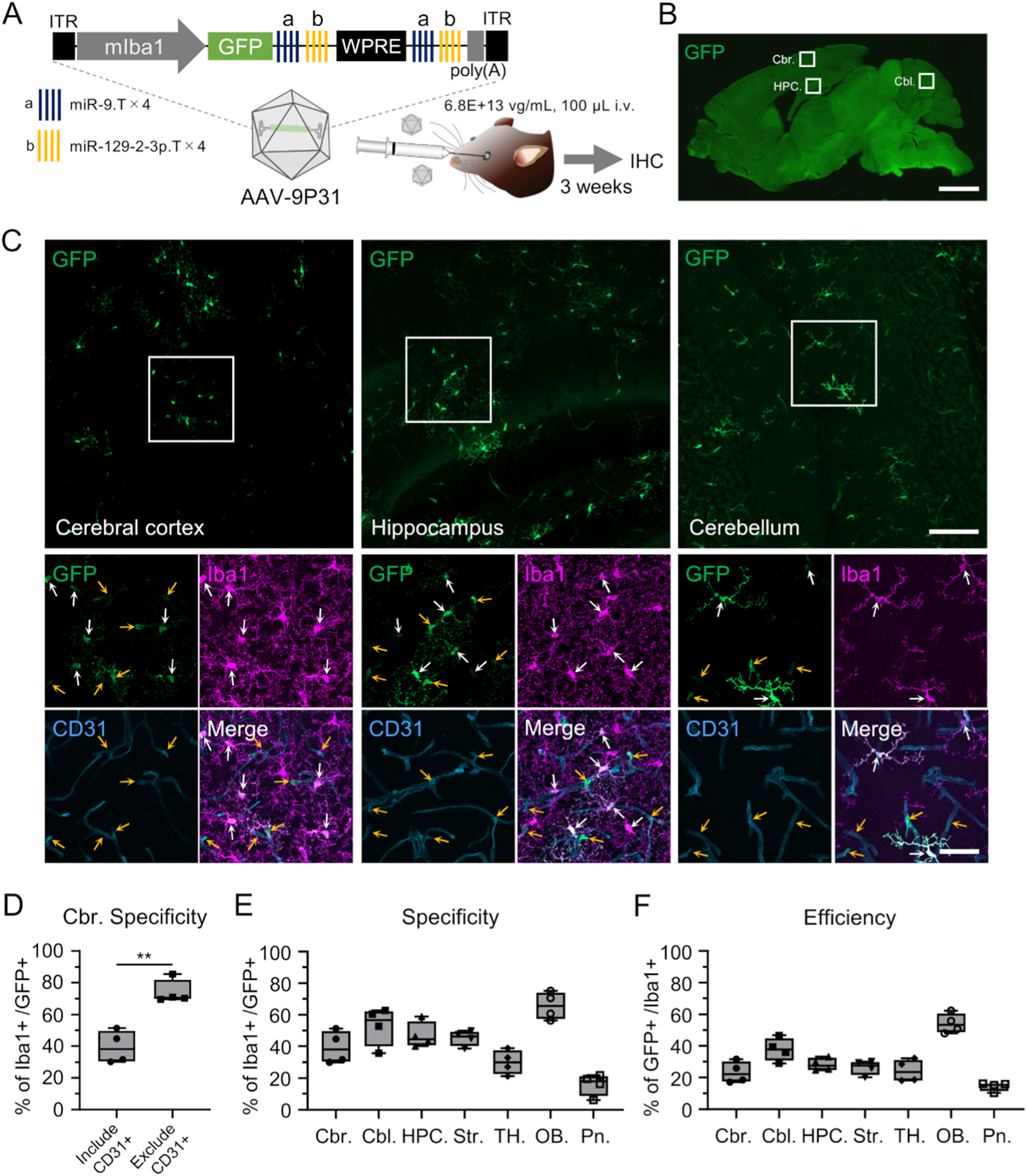
GFP expression in microglia throughout the brain following intravenous injection of AAV-9P31.mIba1.GFP.ab-WPRE-ab. **(A)** The microglia-targeting AAV genome containing mIba1.GFP.ab-WPRE-ab was packaged with the mouse BBB-permeable AAV-9P31 capsid. The AAV vector (6.8E+13 vg/mL, 100 μL) was injected intravenously into C57BL/6J mice via the orbital venous plexus. The mice were euthanized three weeks after injection and analyzed by immunohistochemistry (IHC). **(B)** Sagittal section of the whole brain immunolabeled for GFP. Scale bar, 2 mm. **(C)** The top three images, immunolabeled for GFP, correspond to the boxed areas of the cerebral cortex, hippocampus, and cerebellum in the sagittal section (B). The bottom four images, immunolabeled for GFP, Iba1, and CD31, are magnifications of the boxed areas in the top GFP-labeled images. White and yellow arrows indicate GFP-expressing microglia and non-microglial cells, respectively. Note that most GFP-positive non-microglial cells are CD31- positive vascular endothelial cells. Scale bars: 100 μm (upper right) and 40 μm (lower right). **(D)** Graph depicting the specificity of GFP expression in cortical microglia. “Include CD31+ (same as data in Fig. 8E Cbr.)” and “Exclude CD31+” represent the microglial specificity of GFP expression when calculated including CD31-positive endothelial cells (39.4 ± 8.9%, n = 4 mice) and excluding CD31-positive endothelial cells (74.0 ± 6.6%, n = 4 mice), respectively. **p<0.01 by unpaired t-test. **(E, F)** Specificity (E) and efficiency (F) of GFP expression in microglia across various brain regions following intravenous injection of the microglia-targeting AAV-9P31 vectors. Cbr.: Cerebrum, Cbl.: Cerebellum, HPC: Hippocampus, Str.: Striatum, TH.: Thalamus, OB: Olfactory bulb, Pn: Pons. See also Figure S4-S5.

Confocal microscopy of brain sections from mice that received higher doses of AAV- 9P31 vectors showed numerous GFP-labeled cells throughout the brain (Fig. 8B, C; Fig. S5B- F). Immunohistochemical analysis revealed that the GFP-expressing cells were primarily Iba1- positive microglia and CD31-positive vascular endothelial cells. Quantification of the immunohistochemistry showed that the specificity of GFP expression for microglia in the cerebral cortex was approximately 40% (39.4 ± 8.9%, n = 4 mice). When the analysis was limited to the brain parenchyma, excluding vascular endothelial cells, the specificity of GFP expression for microglia in the cerebral cortex increased to 74% (74.0 ± 6.6%, n = 4 mice, **p < 0.01 by unpaired t-test) (Fig. 8D). The specificity and efficiency of transgene expression for microglia across different brain regions are summarized in Fig. 8E and F, with the olfactory bulb exhibiting the highest specificity and efficiency among the regions examined.

## Discussion

In this study, we demonstrated that relocating miR-9.T and miR-129.T from the 3’ to the 5’ side of WPRE significantly enhances both the specificity and efficiency of transgene expression in cortical microglia. Furthermore, placing miR target sequences on both the 5’ and 3’ sides of WPRE further improved microglial specificity, which remained robust even two months post-AAV injection. Since miRs bind to and cleave their perfectly complementary sequences,^19,20^ the miR target sequences located downstream of WPRE are cleaved, producing GFP mRNA with WPRE, whereas cleavage of upstream miR target sequences results in GFP mRNA alone (Fig. S1B, C). WPRE stabilizes the mRNA,^21^ preventing degradation and facilitating protein translation. In contrast, GFP mRNA lacking both WPRE and poly(A) sequences is rapidly degraded, leading to no GFP protein expression.

Transgene specificity for microglia was notably higher in AAV constructs with miR.T on both sides of WPRE compared to constructs with miR.T only on the 5’ side (Fig. 2D). This increase is likely because the miR.T on the 3’ side acts as a safeguard for the 5’ miR.T. Without the 3’ miR.T, incomplete cleavage at the 5’ miR.T produces mRNA containing GFP, WPRE, and poly(A), resulting in strong GFP protein expression. However, the presence of the 3’ miR.T ensures that any uncleaved mRNA at the 5’ miR.T is cleaved at the 3’ miR.T, producing mRNA consisting only of GFP and WPRE sequences, which significantly suppresses GFP protein expression due to the absence of the poly(A) signal.

Replacing the mIba1 promoter in the microglia-targeting cassette with the CAG promoter resulted in oligodendrocyte-specific transgene expression (Fig. 3), indicating that the CAG promoter is not active in microglia, and the mIba1 promoter is non-functional in oligodendrocytes. AAV.CAG.GFP.ab-WPRE-ab induces mRNA transcription in neurons, astrocytes, and oligodendrocytes. However, mRNA containing miR.T is likely degraded upon binding to endogenous miRs in neurons and astrocytes but remains intact in oligodendrocytes, where miR-9 and miR-129-2-3p are either absent or insufficiently expressed.^22^

Previously, we reported that intravenous injection of AAV.mIba1.GFP.WPRE-ab resulted in residual GFP aggregates in microglial lysosomes.^15^ In this study, we utilized five different BBB-penetrating capsid variants to investigate whether intravenous administration of GFP-expressing AAV with miR.T sequences on both sides of WPRE could specifically and efficiently label microglia. Our results demonstrated that AAV-9P31, AAV-PHP.B, and AAV-PHP.eB expressed GFP in the cytoplasm of microglia, with AAV-9P31 showing the highest efficiency. Further investigation using high doses of AAV-9P31.mIba1.GFP.ab-WPRE-ab revealed microglial specificity ranging from 20% in the pons to 70% in the olfactory bulb, with transduction efficiency ranging from 20% (pons) to 60% (olfactory bulb).

Notably, intravenous administration of AAV-9P31.mIba1.GFP.ab-WPRE-ab resulted in highly efficient GFP expression in vascular endothelial cells across all brain regions examined. These findings suggest that the mIba1 promoter is active in vascular endothelial cells, where miR-9 and miR-129-2-3p are not endogenously expressed, leading to unregulated transgene expression. When transgene expression in vascular endothelial cells was excluded, GFP expression specificity in cortical microglia reached nearly 75% (Fig. 8D), suggesting that intravenous administration of AAV-9P31.mIba1.ab-WPRE-ab enables microglia-selective transgene expression in the brain parenchyma. Incorporating sequences complementary to miRs expressed in vascular endothelial cells but not in microglia into the AAV vector may help suppress transgene expression in endothelial cells and further enhance microglial specificity.

Currently available wild-type AAV serotype capsids tend to have high affinity for neurons but low affinity for microglia,^30,31,32,33,34,35^ resulting in limited transgene expression in microglia. The development of microglia-tropic capsid mutants would enable more specific, efficient, and robust transgene expression in these cells.

Here, by placing miR.T on both sides of WPRE in AAV vectors, we significantly improved the specificity and efficiency of transgene expression in cortical microglia over an extended period. These enhanced microglia-targeting AAV vectors will be valuable tools for studying microglial physiology and pathophysiology, as well as for developing microglia-targeted gene therapies for various neuropsychiatric diseases involving microglia.

## Methods

### Vector construction

The plasmid WPRE-ab (pAAV/mIba1.GFP.WPRE.miR-9.T.miR-129-2-3p.T.SV40pA) was obtained from previous studies in our laboratory (Addgene plasmid #190163). The plasmid CAG-eYFP-3x-miR708-5p-TS was a gift from Viviana Gradinaru (Addgene plasmid #117381; http://n2t.net/addgene:117381; RRID:Addgene_117381). The plasmid pGP-AAV-syn-FLEX-jGCaMP8s-WPRE was a gift from the GENIE Project (Addgene plasmid #162377; http://n2t.net/addgene:162377; RRID:Addgene_162377).

To create the plasmid WPRE-cab (pAAV/mIba1.GFP.WPRE.miR-708-5p.T.miR- 9.T.miR-129-2-3p.T.SV40pA), the miR-708-5p.T from CAG-eYFP-3x-miR708-5p-TS was amplified by PCR and inserted into WPRE-ab at the KpnI restriction enzyme site using the In- Fusion Cloning Kit (TaKaRa Bio, Shiga, Japan). To construct the plasmids ab-WPRE or ab- WPRE-ab (pAAV/mIba1.GFP.miR-9.T.miR-129-2-3p.T.WPRE.SV40pA or pAAV/mIba1.GFP.miR-9.T.miR-129-2-3p.T.WPRE.miR-9.T.miR-129-2-3p.T.SV40pA

[Addgene plasmid #226475]), the miR.Ts were placed either upstream or on both sides of the WPRE in pAAV/mIba1.GFP.WPRE.SV40pA. To create the plasmid pAAV/mIba1.jGCaMP8s.miR-9.T.miR-129-2-3p.T.WPRE.miR-9.T.miR-129-2- 3p.T.SV40pA, the jGCaMP8s gene was amplified by PCR using pGP-AAV-syn-FLEX- jGCaMP8s-WPRE as a template and inserted into ab-WPRE-ab at the AgeI and NotI restriction enzyme sites. The plasmids pAAV/mIba1.GFP.WPRE and pAAV/mIba1.GFP were created by removing the SV40pA or WPRE-SV40pA, respectively, from pAAV/mIba1.GFP.WPRE.SV40pA.

To produce Rep/Cap plasmids for AAV-9P31 or innate-MGs (AAV(H) and AAV(Y)) (Fig. S4), codon-optimized peptide sequences were inserted into the variable region VIII of AAV9. The peptide sequences for innate-MGs were obtained from the patent information in WO 2022/023773 A1.

### AAV vector production and titration

Recombinant single-strand AAV vectors were produced using the ultracentrifugation method as described previously.^36^ In brief, three plasmids—the expression plasmid pAAV, pHelper (Agilent Technologies, Santa Clara, CA, USA), and the Rep/Cap plasmid—were co-transfected into HEK293T cells (HCL4517; Thermo Fisher Scientific, Waltham, MA, USA) using polyethylenimine “Max” (24765-1; Polysciences Inc., Warrington, PA, USA). Cells were cultured in Dulbecco’s Modified Eagle’s Medium (DMEM; D5796- 500ML; Sigma-Aldrich, St. Louis, MO, USA) supplemented with 8% fetal bovine serum (CCP- FBS-BR-500; Cosmo Bio, Tokyo, Japan). Viral particles were harvested from the culture medium 6 days after transfection and concentrated by precipitation with 8% polyethylene glycol 8000 (P5413; Sigma-Aldrich) and 500 mM sodium chloride. The precipitated AAV particles were resuspended in Dulbecco’s phosphate-buffered saline (D-PBS) and purified with iodixanol (Optiprep; AXS-1114542-250ML; Alere Technologies, Oslo, Norway) using linear density gradient ultracentrifugation with an ultracentrifuge (CP80WX; Himac, Tokyo, Japan). The viral solution was further concentrated and formulated in D-PBS using a Vivaspin Turbo 15 MWCO 100000 PES (VS15T42; Sartorius, Göttingen, Germany).

The genomic titers of the purified AAV vectors, except for AAV9.mIba1.GFP, were determined by quantitative real-time PCR (TP900 or TP970; TaKaRa Bio) using Power SYBR Green Master Mix (Thermo Fisher) with primers targeting the WPRE sequence (5′- CTGTTGGGCACTGACAATTC-3′ and 5′-GAAGGGACGTAGCAGAAGGA-3′). For AAV9.mIba1.GFP, which lacks the WPRE sequence, primers targeting GFP (5′- CGACCACTACCAGCAGAACAC-3′ and 5′-TGTGATCGCGCTTCTCGTTGG-3′) were used.

### Animals

Wild-type C57BL/6J mice (6–8 weeks old) were used in this study. Careful attention was given to the sex of the mice to avoid bias. All procedures were performed according to protocols approved by the Japanese Act on the Welfare and Management of Animals and the Guidelines for Proper Conduct of Animal Experiments issued by the Science Council of Japan. The experimental protocol was approved by the Institutional Committee of Gunma University (Nos. 21–065 and 23–057). All efforts were made to minimize suffering and reduce the number of animals used.

### Stereotaxic injection

Mice were anesthetized with intraperitoneal ketamine (80 mg/kg body weight [BW]) and xylazine (1.6 mg/kg BW) and maintained under 0.5% isoflurane anesthesia using an anesthetic vaporizer (MK-AT210; Muromachi Kikai, Fukuoka, Japan). Anesthetic depth was monitored via the toe-pinch reflex. A hole was created over the injection site using a 30G needle. AAV vectors were injected into the bilateral cerebral cortex (M1-M2) and striatum, or into the cerebellar vermis. A 10-μL Hamilton syringe with a 33G needle was used with a stereotaxic micromanipulator (SMM-100; Narishige, Tokyo, Japan) mounted on a stereotactic frame (SRS-5-HT; Narishige). The stereotaxic coordinates relative to bregma were as follows: cerebral cortex (M1-M2) AP −1.0 mm, ML ±1.0 mm, DV +0.8 mm (advanced to +1.0 mm, then retracted by −0.2 mm); striatum AP −1.0 mm, ML ±1.75 mm, DV +2.75 mm; cerebellum AP +6.5 mm, ML 0 mm, DV 1.8 mm (advanced to +2.0 mm, then retracted by −0.2 mm).

### Intravenous injection

Mice were anesthetized by intraperitoneal injection of ketamine (100 mg/kg BW) and xylazine (2.0 mg/kg BW). Anesthetic depth was monitored using the toe-pinch reflex. A 30G needle (08-277; Nipro, Osaka, Japan) was used to access the orbital venous plexus, and 100 μL of AAV solution was slowly injected.

### Immunohistochemistry

At 21 or 56 days post-injection, mice were transcardially perfused with PBS (pH 7.4) followed by 4% paraformaldehyde in 0.1 M phosphate buffer (PB) (pH 7.4). Brains were post-fixed overnight and then transferred to PBS. Brain slices were prepared using a microtome (VT1200S; Leica, Wetzlar, Germany). Coronal slices (50 μm) were used for the cerebral cortex and striatum, and sagittal slices (50 μm) were used for the cerebellum and whole brain. Slices were incubated with primary antibodies (Table S1) in a blocking solution (2% normal donkey serum, 2% BSA, 0.5% Triton X-100, and 0.05% NaN₃ in 0.1 M PB) overnight at 4 °C. After six washes with PBS, slices were incubated with secondary antibodies (Table S1) in a blocking solution for 3 hours at room temperature (24–26 °C). Slices were then washed six times with PBS and mounted on glass slides with ProLong Diamond Antifade Mountant (Thermo Fisher Scientific). In Fig. 3, staining was performed with two antibodies per slice due to limitations in the number of wavelengths observable in our setup.

### Imaging analysis

The sagittal sections of the whole brain from intravenously injected mice were acquired using a fluorescence microscope (BZ-X800; Keyence, Osaka, Japan). All other immunostained slices were imaged using a confocal laser-scanning microscope (LSM 800; Carl Zeiss, Oberkochen, Germany). The number of cells was manually counted using z-stack images with ZEISS ZEN software (ZEN 3.6, Carl Zeiss). GFP fluorescence intensity was measured using ImageJ software (https://imagej.net/software/fiji/downloads).

### Confocal live imaging in microglia of the motor cortex

Confocal live Ca²⁺ or GFP imaging of microglia was performed in acute cerebral slices from the mouse motor cortex, including primary and secondary motor areas, as described previously,^15^ with some modifications. Coronal slices (250–300 µm thick) of the cerebral cortex containing the motor areas were prepared using a vibroslicer (VT1200S; Leica, Germany) 3– 4 weeks after AAV injection into the motor cortex (0.5–5 µL at a titer of 0.4–5 × 10¹³ vg/mL for jGCaMP8s; 1 µL at a titer of 0.1 × 10¹³ vg/mL for GFP). The slices were maintained in artificial cerebrospinal fluid (ACSF) containing (in mM): 125 NaCl, 2.5 KCl, 2 CaCl₂, 1 MgCl₂, 1.25 NaH₂PO₄, 26 NaHCO₃, and 10 D-glucose, bubbled with 95% O₂ and 5% CO₂ at room temperature for over one hour before recording. Image acquisition, processing, and analysis were performed using Andor iQ3 (Andor), NIH ImageJ, Igor Pro9 (WaveMetrics) with Neuromatic (http://www.neuromatic.thinkrandom.com),^37^ and custom-written programs by NH.

The ‘jGCaMP8s’ sensor protein is a fast and sensitive genetically encoded Ca²⁺ indicator that increases fluorescence upon an elevation in intracellular Ca²⁺ concentration.^23^ To monitor Ca²⁺ signals in jGCaMP8s-expressing microglia in brain slices, confocal fluorescence images were acquired every 2 seconds using a water-cooled EM-CCD camera (200 ms exposure time, 512 × 512 pixels; iXon3 DU-897E-CS0-#BV-500; Andor, Belfast, Northern Ireland), a 40× water immersion objective (LUMPLFLN 40XW; Olympus, Tokyo, Japan), and a high-speed spinning-disk confocal unit (CSU-X1; Yokogawa Electric, Tokyo, Japan) attached to an upright microscope (BX51WI; Olympus, Tokyo, Japan). A blue laser light (488 nm; Stradus 488-50; VORTRAN, Sacramento, CA) was used for excitation, and emitted fluorescence was collected through a 500–550 nm band-pass filter. During recordings, cortical slices were continuously perfused with ACSF bath solution at room temperature. ATP (100 µM) dissolved in ACSF was bath-applied for 1.5–4 minutes via a gravity-fed bath-application device to elicit intracellular Ca²⁺ increases and process movement in microglia. Image drift (translation drift) during recordings was corrected using the Image Stabilizer plugin (http://www.cs.cmu.edu/~kangli/code/Image_Stabilizer.html) or the Template Matching and Slice Alignment plugin (https://sites.google.com/site/qingzongtseng/template-matching-ij-plugin#downloads) in ImageJ.

GCaMP fluorescence intensity at time t (Fₜ) in each pixel was background-subtracted, and Ca²⁺-dependent relative changes in fluorescence were measured by calculating ΔF/Fbasal, where Fbasal is the basal fluorescence intensity averaged during pre-stimulus frames (i.e., more than 10 frames before ATP application) and ΔF = Fₜ - Fbasal. Background fluorescence was measured from regions devoid of cellular structures in the same frame. Mean ΔF/Fbasal values were calculated for each region of interest (ROI) in each frame. ROIs were placed on GCaMP-positive cellular structures. Because we could not determine whether multiple neighboring ROIs in a frame corresponded to the same cell, each ROI was referred to as a microglial cellular compartment. Ca²⁺ imaging data were collected from 15 different fields of view, each separated by more than 200 µm, in eight slices from five mice. Considering the coverage of microglial process territories (convex hull areas ranging from 800 to 3800 µm², corresponding to circles with diameters of several tens of micrometers),^26^ we estimate that the Ca²⁺ imaging data originated from more than 15 microglia. To quantify ATP-induced Ca²⁺ signals in GCaMP-positive cells, the peak amplitude of ΔF/Fbasal was measured within a 100-second time window after the onset of ATP application.

To capture live morphological dynamics in GFP-positive microglia, confocal GFP fluorescence images were acquired every 2 seconds under the same camera settings as for GCaMP imaging (single focal plane imaging). This method often missed out-of-focus microglial processes (Fig. 7, Single focal plane) due to the complex three-dimensional structure of microglia.^26^ To overcome this limitation, z-stack images were acquired at multiple focal planes, covering the entire microglial structure with a depth range of 40–73 µm. These z-stacks were obtained at each time point every 6–9 seconds through rapid z-axis positioning of the objective lens (z-step size: 1.65–1.94 µm) using a piezo objective scanner (PFM450E, Thorlabs). Maximum intensity projections of the z-stacks on the xy plane were generated to create 2D time-lapse images. The signal-to-noise ratio of the GFP signal was significantly higher than that of the GCaMP signal under these experimental conditions. Image drift during GFP imaging was corrected using the same method as in Ca²⁺ imaging.

### Statistics

GraphPad PRISM (version 9; GraphPad Software, San Diego, CA, USA) was used for statistical analyses and data visualization. Specificity, efficiency, and fluorescence intensity data in Fig. 2F were presented as box-and-whisker plots. The box-and-whisker plots depict the median (centerlines), 25th and 75th percentiles (bounds of the box), and minimum/maximum values (whiskers). Statistical significance for specificity and efficiency was determined using unpaired t-tests or Bonferroni’s multiple comparisons test after one-way ANOVA. Statistical significance for fluorescence intensity was determined using unpaired t-tests (Fig. 2F). Comparisons of fluorescence intensity at 3 weeks versus 2 months post-injection were presented as cumulative plot distributions and analyzed with the Kolmogorov-Smirnov test (Fig. 5E, F). Unless otherwise indicated, data are presented as mean ± standard deviation (SD) in the Results text or Figure legends.

### Data availability

All relevant data related to this manuscript will be made available upon reasonable request to the corresponding authors. Detailed information on pAAV/mIba1.GFP.miR-9.T.miR-129-2- 3p.T.WPRE.miR-9.T.miR-129-2-3p.T.SV40pA, including its sequence, can be obtained from Addgene (Addgene plasmid #226475).

## Acknowledgements

This work was supported by grants from the Program for Brain Mapping by Integrated Neurotechnologies for Disease Studies (Brain/MINDS; JP20dm0207057/JP21dm0207111 [to H.H.]) and Multidisciplinary Frontier Brain and Neuroscience Discoveries (Brain/MINDS 2.0; JP24wm0625103 [to H.H.]) from the Japan Agency for Medical Research and Development (AMED), and MEXT/JSPS KAKENHI (20K06906/24K10022 [to N.H.], 22K06454/24H01221 [to A.K.] and 23H02791 [to H.H.]). The authors thank Asako Ohnishi, Nobue McCullough, and Chieko Miyazawa for AAV vector production; Junko Sugiyama for mouse care; Syota Togai for conducting the preliminary CAG experiments; and Yasunori Matsuzaki and Yuki Fukai for their technical advice on plasmid editing and animal experiments. The graphical abstract includes illustrations created with BioRender.com.

## Author contributions

R.A., A.K., and H.H. designed the experiments; R.A., N.H., and H.K. conducted the research; N.H. performed the Ca²⁺ imaging and live microglia imaging experiments; H.H. supervised and completed the study.

## Supplemental information

Document S1. Figures S1-S5 and Table S1. Video S1. Source video data, related to Figure 6.

Video S2-S9. Source video data, related to Figure 7.

