## Supplementary material for "Development of an AAV Vector System for Highly Specific and Efficient Gene Expression in Microglia": Document S1

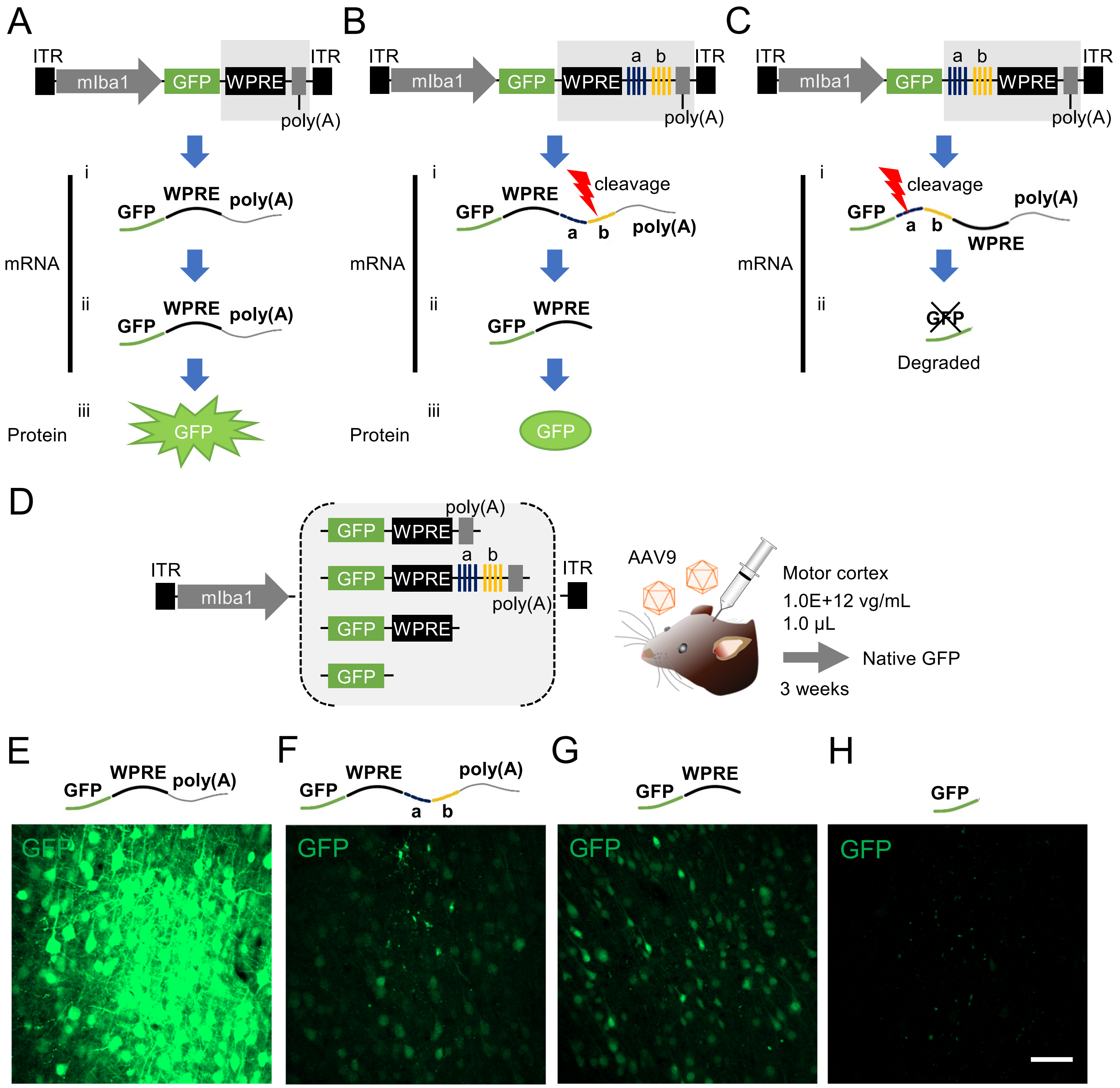


**Figure S1. A hypothetical model and experiments demonstrating that GFP expression is suppressed when miR.T is located on the 5' side of WPRE, related to Figure 2.**

**(A)** AAV.mIba1.GFP.WPRE.poly(A) produces mRNA consisting of GFP, WPRE, and poly(A) signal, resulting in strong GFP expression. **(B)** Insertion of miR.Ts on the 3' side of WPRE produces mRNA containing GFP, WPRE, miR.Ts, and poly(A) signal. This mRNA is cleaved at the miR.Ts by RNA interference in neurons. The processed GFP and WPRE mRNA may still be translated into GFP, although less efficiently than without miR.Ts (as shown in (A)). **(C)** Insertion of miR.Ts on the 5' side of WPRE produces mRNA containing GFP, miR.Ts, WPRE, and poly(A) signal. This mRNA is likely processed into mRNA containing only GFP, which may lead to its degradation. **(D)** Experimental design to test this hypothesis. AAV.mIba1.GFP with or without WPRE, miR.T, and poly(A) signal was prepared as indicated. Mice received cerebral injections of one of the AAVs and were euthanized three weeks later to observe native GFP expression. **(E-H)** Confocal micrographs of cerebral sections from mice injected with AAVs as shown in (D). The mRNA transcribed from the injected AAV is illustrated above each panel. Scale bar: 50 µm.

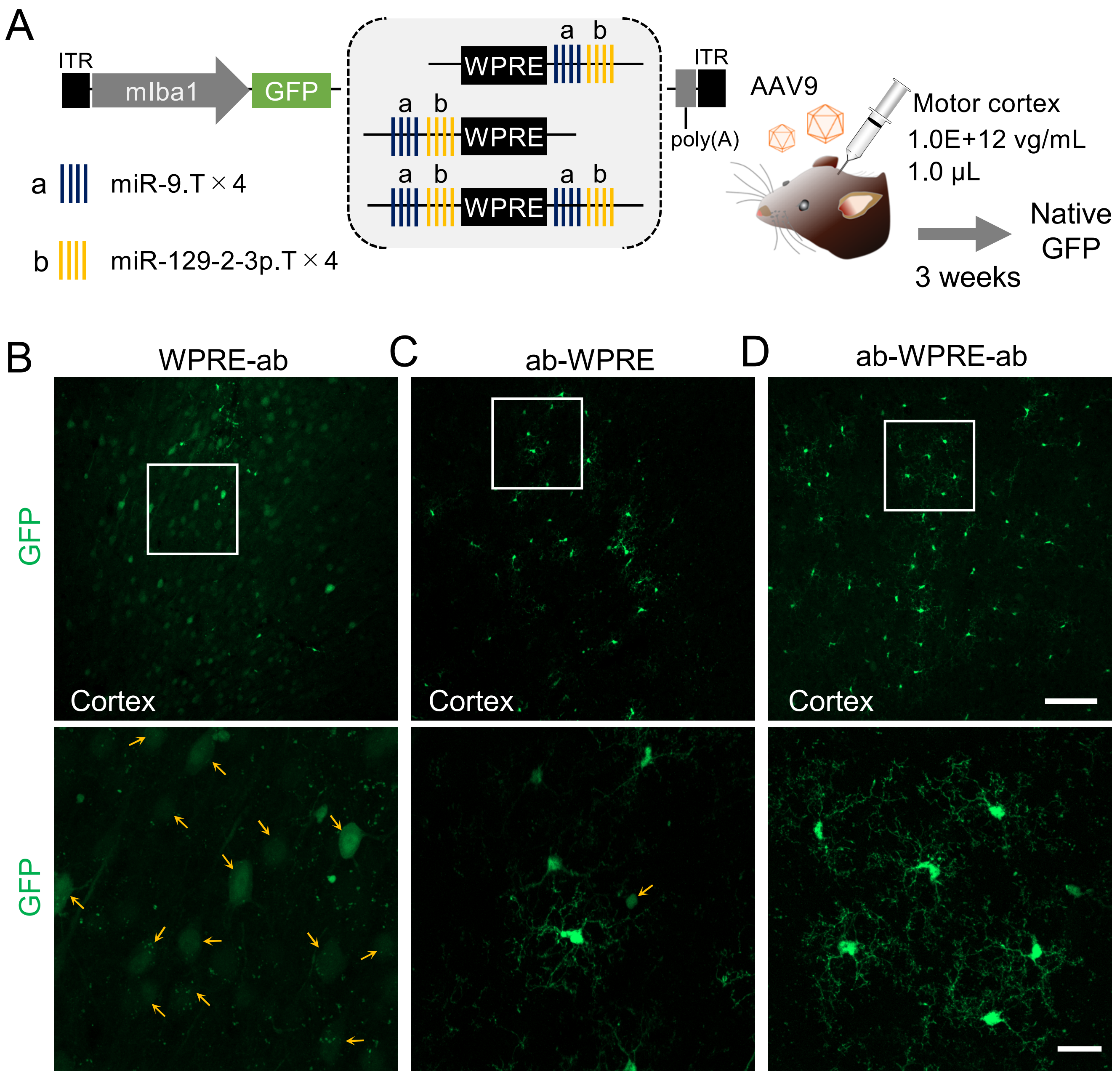


**Figure S2. Native GFP fluorescence in cortical microglia of mice injected with AAV9.mIba1.GFP.WPRE, with miR.T on the 3', 5', or both sides of WPRE, related to Figure 2.**
**(A)** Schematic showing three AAV genomes with miR.T sequences positioned downstream, upstream, or on both sides of WPRE. One of these AAVs was injected into the motor cortex at the indicated doses, and the cerebral cortex was sectioned three weeks after injection to observe native GFP fluorescence. **(B-D)** Native GFP fluorescence images of the cerebral cortex. The boxed areas in the lower magnification images are enlarged and shown below. Yellow arrows indicate cells morphologically distinct from microglia. Scale bars: 100 µm (upper right) and 20 µm (lower right).

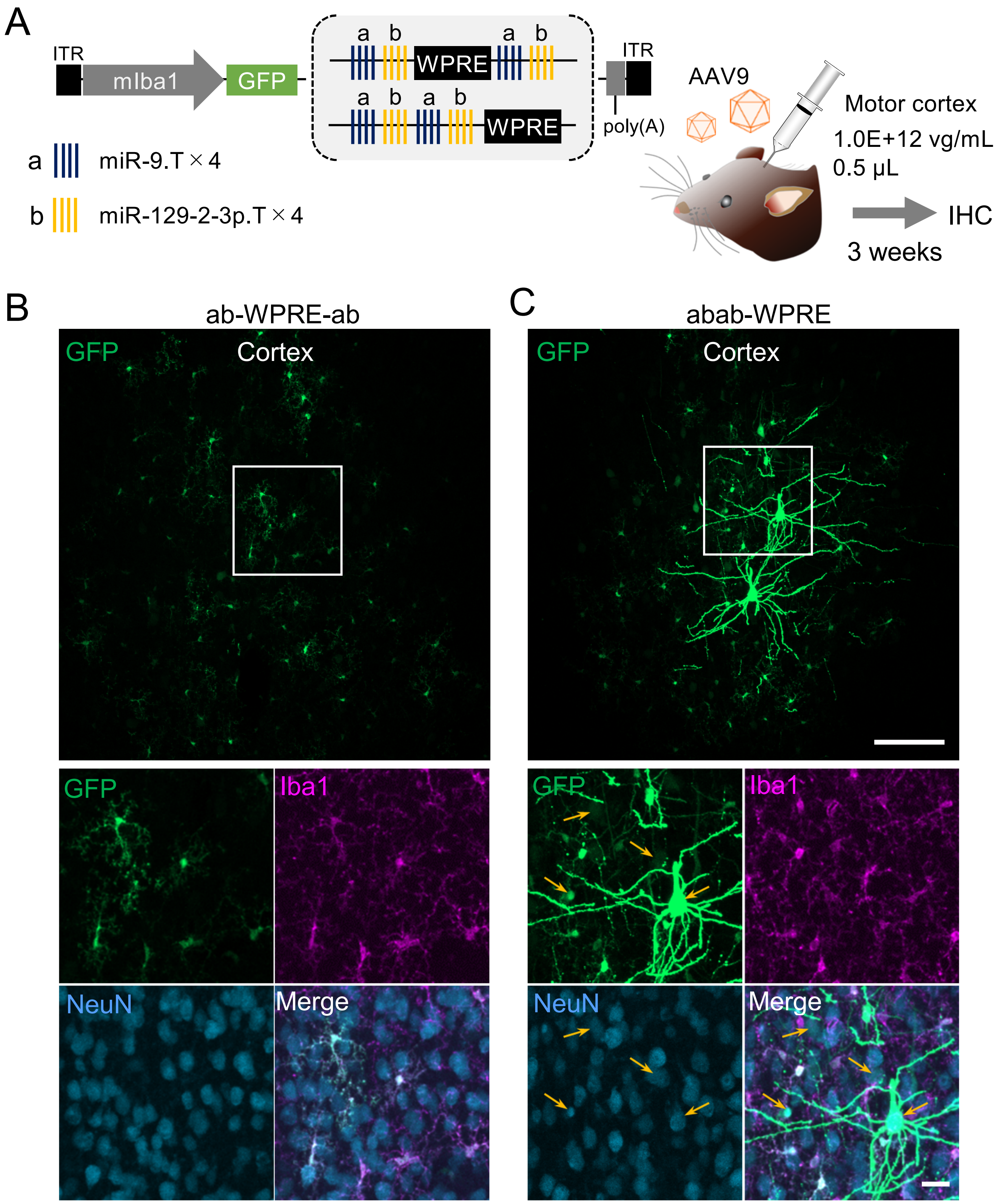


**Figure S3. Placing miR.Ts on both sides of the WPRE is crucial for increasing the specificity of GFP expression in microglia, related to Figure 2.**

**(A)** AAV.mIba1.GFP with two sets of miR.Ts on each side of WPRE, or with four sets of miR.Ts only upstream of WPRE, was injected into the mouse cerebral cortex at the indicated doses. **(B, C)** Confocal microscopy of cortical sections three weeks after injection of AAV.mIba1.GFP with ab-WPRE-ab (B) or abab-WPRE (C). The boxed areas in the low-magnification images are shown enlarged below. Yellow arrows indicate NeuN- and GFP-double positive neurons. Scale bars: 100 μm (upper right) and 20 μm (bottom right).

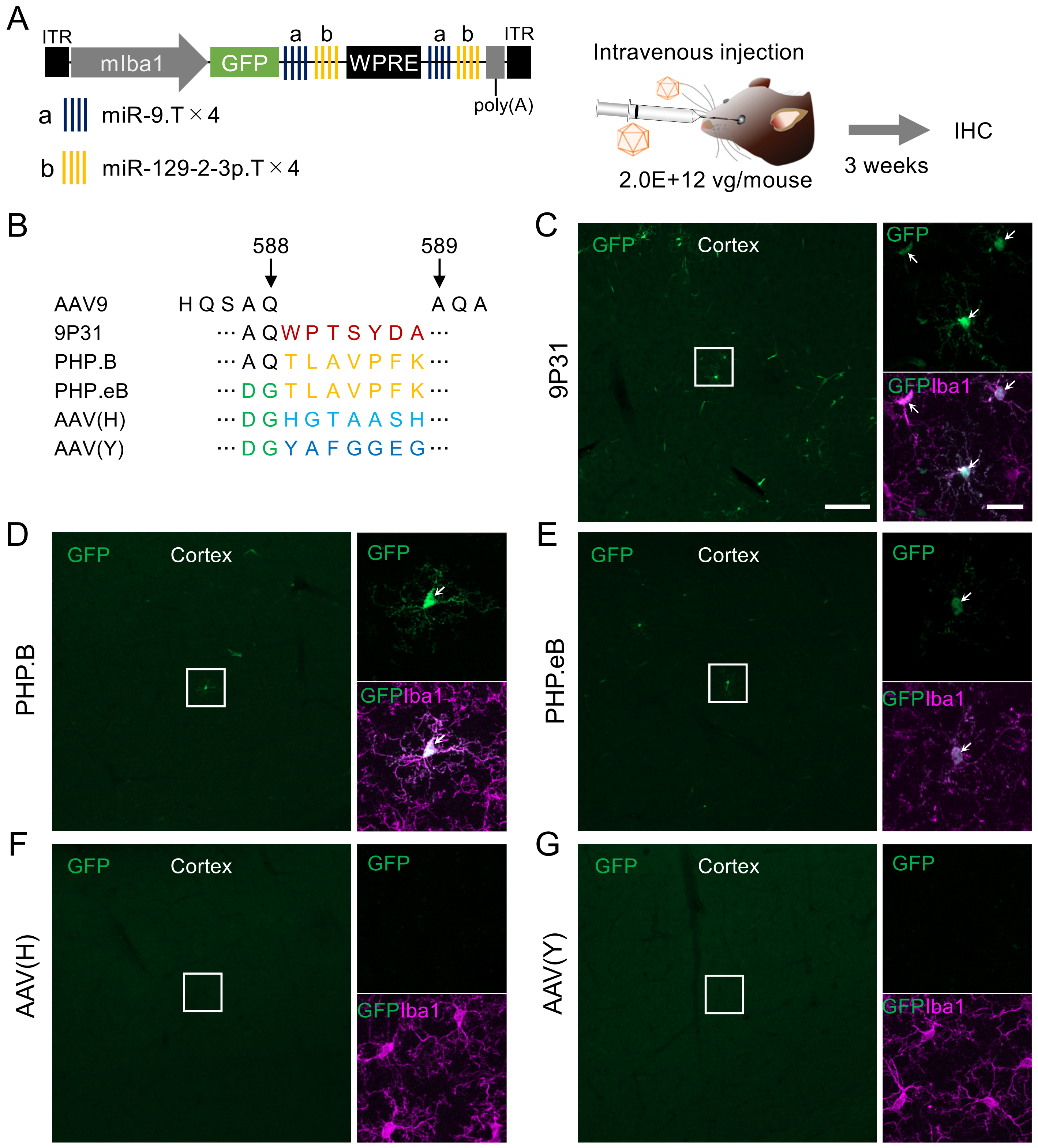


**Figure S4. Comparison of five BBB-permeable capsid mutants for gene expression in microglia upon intravenous administration, related to Figure 8.**

**(A)** Five different BBB-penetrating capsid vectors containing mIba1.GFP.ab-WPRE-ab.poly(A) were injected intravenously into mice via the orbital plexus. **(B)** The seven amino acid insertions between 588Q and 589A of the AAV9 capsid in the BBB-penetrating capsid variants (including additional flanking mutations) are presented alongside the parental AAV9 sequence. **(C–G)** Confocal microscopy images of cortical sections from mice injected intravenously with microglia-targeting, BBB-permeable AAV vectors. The BBB-penetrating capsid mutants used are shown at the left side of each panel. Boxed areas are enlarged and displayed on the right sides. Arrows indicate microglia double immunostained for GFP and Iba1. Scale bars in panel (C): 100 µm (left) and 20 µm (lower right).

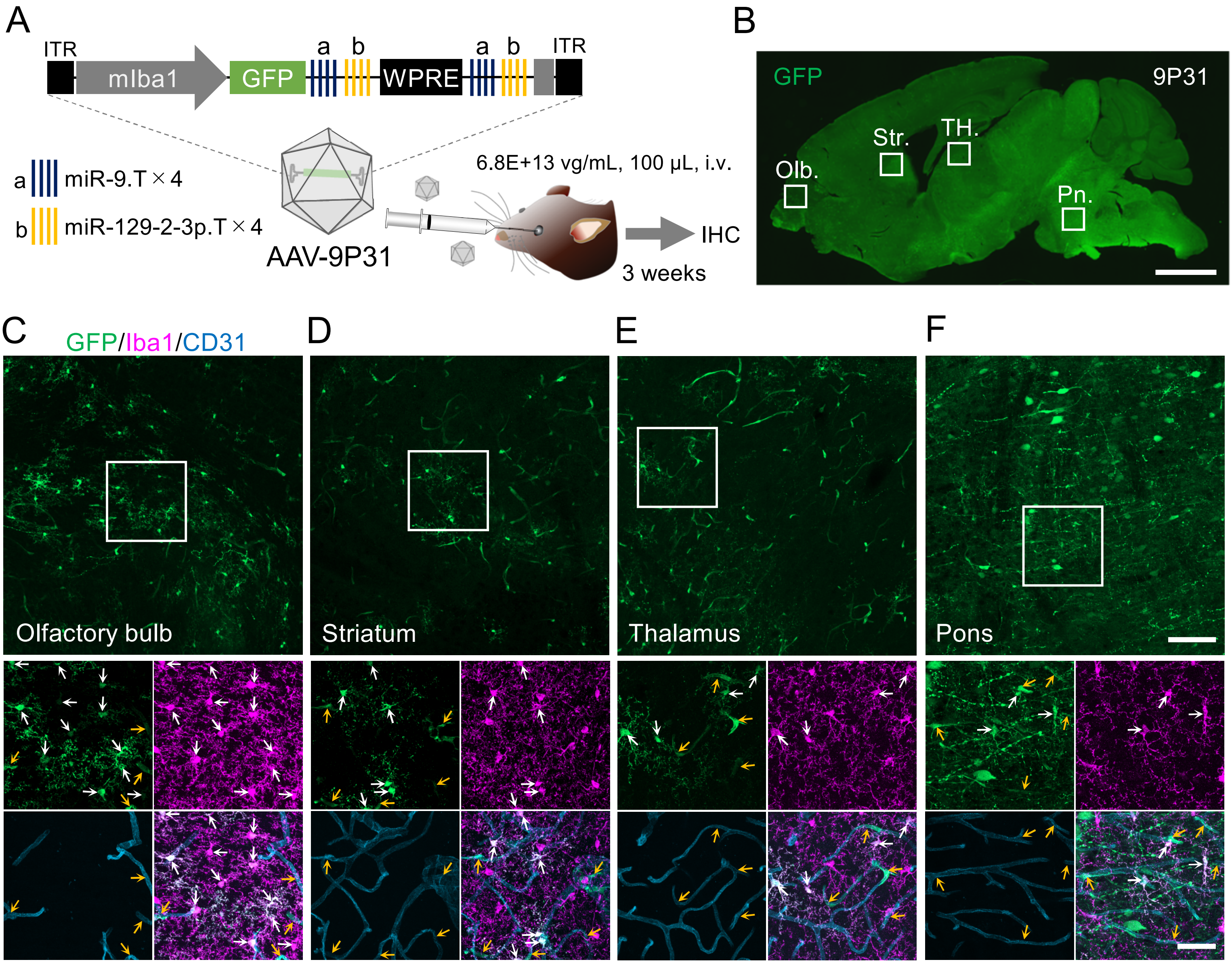


**Figure S5. GFP expression in microglia and brain microvascular endothelial cells across various brain regions following intravenous injection of microglia-targeting AAV-9P31 vectors, related to Figure 8.** **(A)** High dose of AAV-9P31.mIba1.GFP.ab-WPRE-ab.poly(A) (6.8E+13 vg/mL, 100 µL) was intravenously injected. Sagittal brain sections were prepared, and GFP-expressing cell types were examined by immunohistochemistry (IHC). **(B)** GFP immunolabeling image of a sagittal brain section from a mouse intravenously injected with AAV-9P31. **(C–F)** Upper panels: Enlarged GFP immunofluorescent images of the boxed regions in (B): thalamus (C), olfactory bulb (D), striatum (E), and pons (F). Lower panels: Further enlarged immunohistochemical images of the boxed areas in the respective upper panels. Microglia and vascular endothelial cells were immunolabeled with anti-Iba1 (magenta) and anti-CD31 (blue) antibodies, respectively. White and yellow arrows indicate GFP-positive microglia and vascular endothelial cells, respectively. Scale bars in the upper and lower panels of (F): 100 µm and 40 µm, respectively.

**Table S1. Antibody list**

| Primary antibody | | | | | | |
|---|---|---|---|---|---|---|
| No. | Antibody | Host | Monoclonal/ Polyclonal | Dilution ratio | Source | Identifier |
| 1 | anti-GFP | Rat | Mono | ×1000 | Nacalai | 04404-84 |
| 2 | anti-Iba1 | Rabbit | Poly | ×500 | Wako | 019-19741 |
| 3 | anti-NeuN | Mouse | Mono | ×1000 | Chemicon | MAB377 |
| 4 | anti-S100β | Rabbit | Poly | ×200 | Frontier Institute | S100β-Rb-Af1000 |
| 5 | anti-Olig2 | Mouse | Mono | ×500 | Sigma | MABN50 |
| 6 | anti-GFP | Goat | Poly | ×200 | Frontier Institute | GFP-Go-Af1480 |
| 7 | anti-CD31 | Rat | Poly | ×100 | BD pharmingen | 550274 |

| Secondary antibody (Alexa Fluor Plus) | | | | | | | |
|---|---|---|---|---|---|---|---|
| No. | Antibody | Wavelength | Host | Monoclonal/ Polyclonal | Dilution ratio | Source | Identifier |
| 1 | anti-Rat IgG | 488 | Donkey | Poly | ×2000 | Thermo Fisher | A48269 |
| 2 | anti-Rat IgG | 647 | Donkey | Poly | ×2000 | Thermo Fisher | A48272 |
| 3 | anti-Rabbit IgG | 555 | Donkey | Poly | ×2000 | Thermo Fisher | A32794 |
| 4 | anti-Mouse IgG | 647 | Donkey | Poly | ×2000 | Thermo Fisher | A32787 |
| 5 | anti-Goat IgG | 488 | Donkey | Poly | ×2000 | Thermo Fisher | A32814 |

| Antibody used in each Figure | | |
|---|---|---|
| Figure | Primary | Secondary |
| Fig. 1B, C | 1, 2, 3 | 1, 3, 4 |
| Fig. 2B, C | 1, 2, 3 | 1, 3, 4 |
| Fig. 3B | 1, 2, 3 | 1, 3, 4 |
| Fig. 3C (Iba1, NeuN) | 1, 2, 3 | 1, 3, 4 |
| Fig, 3C (S100β, Olig2) | 1, 4, 5 | 1, 3, 4 |
| Fig. 4B, C | 1, 2, 3 | 1, 3, 4 |
| Fig. 5B, C | 1, 2, 3 | 1, 3, 4 |
| Fig. 8B, C | 2, 6, 7 | 2, 3, 5 |
| Fig. 8D (Cell count for Exclude CD31+) | 2, 6, 7 | 2, 3, 5 |
| Fig. 8E, F (Cell count) | 1, 2, 3 | 1, 3, 4 |
| Fig. S3B, C | 1, 2, 3 | 1, 3, 4 |
| Fig. S4C-G | 1, 2, 3 | 1, 3, 4 |
| Fig. S5B-F | 2, 6, 7 | 2, 3, 5 |

**Video S1. Source video data, related to Figure 6**

This video shows GCaMP images illustrating microglial process movements and ATP-induced $Ca^{2+}$ response as depicted in Fig. 6C. Exogenous ATP (100 μM) was bath-applied for 1.5 minutes, as indicated in the video. The video is played at 30 frames per second (60 times faster than real time). Scale bar: 10 μm.

**Videos S2-5. Source video data, related to Figure 7**

These videos show examples of basal microglial process movements from different microglia in the motor cortex. Supplementary Videos 2 and 3 display time-lapse GFP images captured at a single focal plane, while Supplementary Video 4 and 5 present 2D time-lapse GFP images generated from z-axis maximum intensity projections across multiple focal planes. Notably, putative microglial cell bodies near the center remained static in the projected videos. The videos are played at 60 frames per second (120 times faster than real time) in Supplementary Videos 2 and 3, and at 15 frames per second (100 and 120 times faster than real time) in Supplementary Videos 4 and 5, respectively. Scale bar: 10 μm.

**Video S6. Source video data, related to Figure 7**

This video of GFP images shows ATP-induced microglial process extension (single focal plane) as depicted in Fig. 7B (upper panel). The extended microglial processes reached the focal plane and became visible with a slight delay following bath application of ATP. The video is played at 60 frames per second (120 times faster than real time). Scale bar: 10 μm.

**Videos S7-9. Source video data, related to Figure 7**

These videos are 2D time-lapse GFP images constructed from the maximum intensity projections of z stacks acquired from putative single microglia. The videos demonstrate ATP-induced extension and subsequent increased motility of microglial processes in the motor cortex. They are played at 30 frames per second (180 times faster than real time). Scale bar: 10 μm.
